# Enhancing cortico-motoneuronal projections for vocalization in mice

**DOI:** 10.1101/2024.10.14.618267

**Authors:** J. Lomax Boyd, Louisa Kuper, Elena Waidmann, Victor Yang, Erich D. Jarvis

## Abstract

Several hypotheses have been proposed on the anatomical brain differences that endow some species with the rare ability of vocal learning, a critical component of spoken language. One long-standing thus far untested hypothesis is that a robust direct projection from motor cortex layer 5 neurons to brainstem vocal motor neurons enables fine motor control of laryngeal musculature in vocal learners. This connection has been proposed to form from specialized expression of axon guidance genes in human speech layer 5 neurons and the equivalent songbird neurons of the robust nucleus of the arcopallium. Here we generated mice with conditional knockdown of an axon-guidance receptor, *PLXNA1,* in motor cortex layer 5 neurons, to recapitulate the human and songbird brain expression patterns. These mice showed enhanced layer 5 cortical projections to brainstem vocal motor neurons, increased functional connectivity to phonatory muscles, and displayed a wider range of vocal abilities depending on developmental and social contexts. Our findings are consistent with the theory that direct vocal cortico-motoneuronal projections influence vocal behaviors.

## Introduction

Vocal production learning, the ability to produce novel sounds through vocal imitation^1^ (herein vocal learning), is a highly derived and rare trait essential for spoken language in humans^2^. Advanced vocal learning has evolved independently in five groups of mammals (humans, cetaceans, pinnipeds, bats, and elephants) and three groups of birds (songbirds, parrots, and hummingbirds). It is distinguished from auditory learning, the ability to make sound associations, and vocal usage learning, the ability to learn to produce innate (or learned) vocalizations in different contexts, both behaviors more ubiquitous among the animal kingdom^1,2^.

To explain the differences between vocal learners and their closest vocal non-learning relatives, a number of anatomical specializations and mechanisms have been proposed (reviewed in^2^). These include: increased brain size or neuron density allowing more space for vocal learning and speech circuits to form; changes in vocal organ anatomy allowing greater variety of vocalizations; duplication and neofunctionalization of forebrain motor learning pathways for vocal learning; specialized direct connections from the auditory to vocal motor areas of the cortex; or specialized direct projections from motor cortex-to-brainstem vocal motor neurons. The later hypothesis, coined the Kuypers-Jürgens hypothesis by Fitch and colleagues^3^, argues that wherever direct motor cortex to brainstem motor neurons have been found in species, it is correlated with greater fine motor control of the associated muscle groups that gives rise to a behavior, in this case, vocal learning. In all vocal learners where this connection has been studied to date (humans, bats, songbirds, parrots, and hummingbirds), they have been found to have a robust direct connection to brainstem vocal motor neurons, whereas close relatives do not^4–6^.

Our group previously discovered that like vocal learners, mice have a motor cortex region active during vocalization (putative laryngeal motor cortex, LMC), that has layer 5 neuron projections to the brainstem vocal motor neurons (nucleus ambiguus, Amb)^7,8^. However, unlike in vocal learners, the projection in mice is much sparser, and the vocal connected layer 5 neurons are intermingled with projection neurons that perform other functions. Further, in vocal learners (humans and the vocal learning birds), these cortical projection neurons were found to have convergent specialized regulation of several hundred genes, including down regulation of repulsive axon guidance ligands or receptors^9–11^. This down-regulation was not found in the mouse motor cortex region that innervates Amb. These and other findings led to the continuum hypothesis of vocal learning, where the degree of the direct projection correlates with the degree of vocal complexity, with mice having a rudimentary degree^2,7,12,13^. It was further hypothesized that the downregulation of repulsive axon guidance receptors or ligands in speech and song motor cortex layer 5 neurons, functionally inhibited repulsion to brainstem vocal motor neurons, allowing cortical axons to form direct monosynaptic connections to them (**Fig. 1a**)^2,10^. A recent study found that one axon guidance receptor known to be lower in layer 5 neurons of human motor cortex, the *PLXNA1* receptor, when downregulated in all layers throughout mouse cortex resulted in increased direct innervation of forelimb motor neurons in the spinal cord and acquisition of greater control over highly dexterous, fine motor movements^14^. This hypothesis demonstrates that loss of gene function can be associated with a gain-of-function phenotype characterized by connectivity and behavior.

**Fig. 1.**
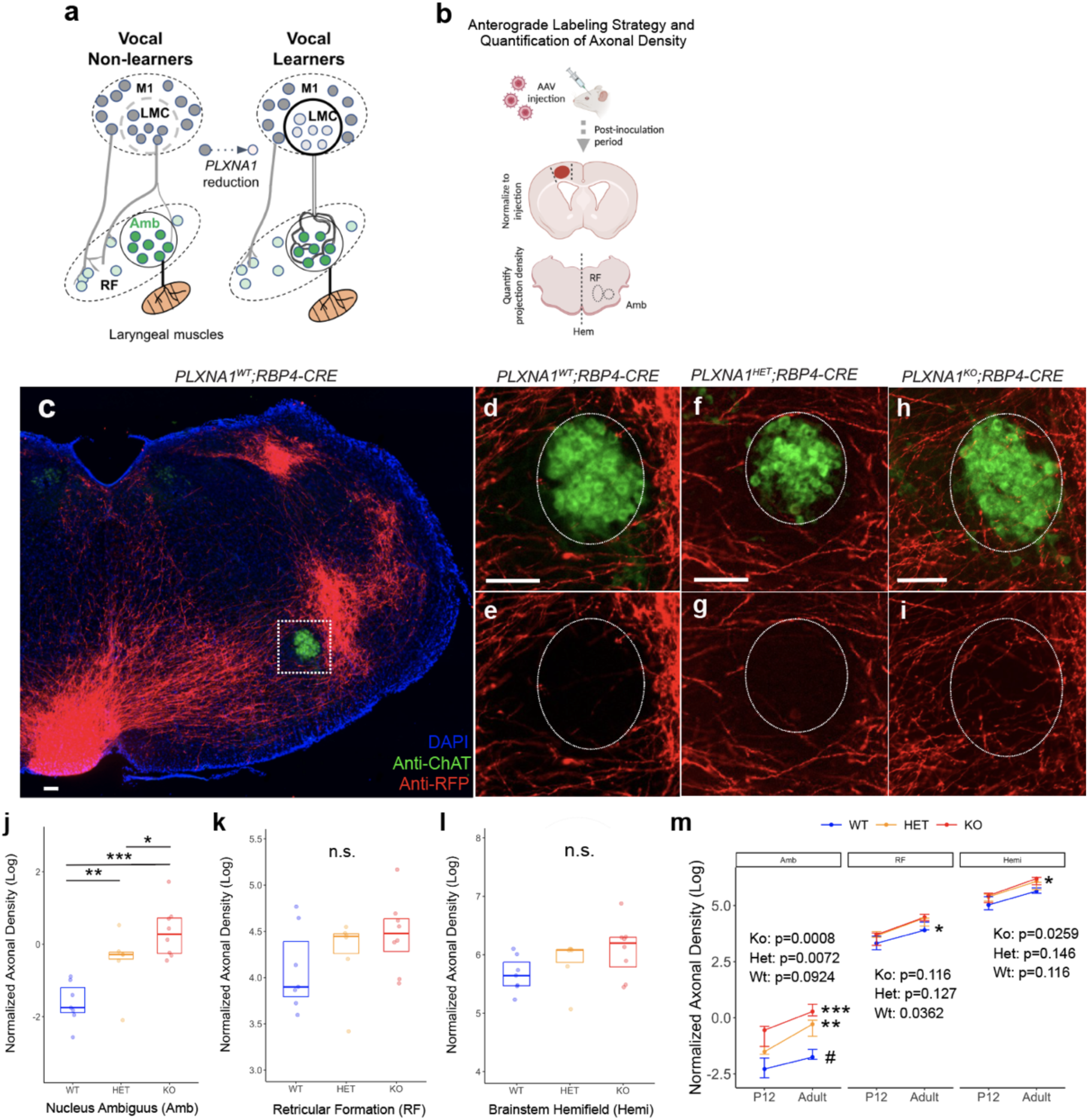
Anterograde tracing of vocal cortico-motoneuronal projections in adult mice. **a**, Proposed repulsion hypothesis mechanism of how direct layer 5 cortical projections to vocal motor neurons are formed in vocal learning species, using *PLXNA1* as an example receptor expressed in pre-synaptic cortical neurons. Potential ligands are *SEMA* and *SLIT* molecules expressed in post-synaptic Amb motor neurons. **b,** Illustration of AAV1.syn.tdTomato injected into primary motor cortex (M1)/laryngeal motor cortex (LMC) region in conditional knockout adult mice and quantification axonal density (N_WT_=7, N_HET_=6, N_KO_=8, mice injected). **c**-**e,** tdTomato+ (red) cortical projections in wildtype mice are found throughout the brainstem with reduced innervation in the vicinity of ChAT+ (green) motor neurons in the Amb (dotted box) **d-i**, Notable increases in axonal fibers are found in *PLXNA1^HET^;RBP4-CRE* and *PLXNA1^KO^;RBP4-CRE* mice relative to wildtype controls. **j**-**l,** Quantification of tdTomato+ (red) axonal fibers, normalized to cortical injection area, in the nucleus ambiguus (Amb), reticular formation (RF), and brainstem hemifield (hemi), contralateral to the side of injection. Axonal density significantly higher in *PLXNA1^KO^;RBP4-CRE* compared to *PLXNA1^WT^;RBP4-CRE* (p<0001, t=-7.06, df=28.8, n=6-8 mice per genotype) and *PLXNA1^HET^;RBP4-CRE* (p=0.0249, t=-2.78, df=28.8) animals; and in *PLXNA1^HET^;RBP4-CRE* compared to *PLXNA1^WT^;RBP4-CRE* controls (p=0.0016, t=-3.87, df=28.8). Boxplots visualize the 75^th^ upper and lower 25^th^ quartile of data with horizontal lines indicating median values. **m**, Comparison of Amb axonal density between P12 pups (**Supplementary Fig. 2**) and adult mice revealed significant developmental increases for *PLXNA1^KO^;RBP4-CRE* and *PLXNA1^HET^;RBP4-CRE* mutants with marginal increases for *PLXNA1^WT^;RBP4-CRE* controls. Points represent the median axonal density with bar representing standard error (s.e). Tukey adjusted p-value for multiple test correction. Scale bar, 100µm. ‘***’ p<0.001, ‘**’ p<0.01, ‘*’ p<0.05, ‘_#_’ p<0.1, ‘n.s.’ non-significant.

In this study, we set out to experimentally test the Kuypers-Jürgens hypothesis of direct vocal cortico-motoneuron connections. Changes in vocal complexity would support the hypothesis; no changes or simplification of vocal plasticity, coordination of features, or syntax would reject the hypothesis. We created transgenic conditional knockout mice with *PLXNA1* expression reduced in layer 5 cortical neurons. These mice showed increased direct innervation of Amb from LMC layer 5 neurons, faster timing to activate vocal motor neurons after stimulation, and a greater complexity of vocal acoustic and temporal features, depending on social context.

## Results

### *PLXNA1* is downregulated in songbird and human song and speech motor cortex

Although *PLXNA1* was among the convergently downregulated axon guidance genes detected by RNA-Seq and microarrays in the songbird and human vocal production cortical areas, validation studies were not followed up on this gene^11^. We examined *in-situ* hybridization expression data available from the Zebra Finch Expression Brain Atlas (ZEBrA)^15^ and confirmed *PLXNA1* down-regulation in the robust nucleus of the arcopallium (RA) song nucleus, which contain projection neurons to phonatory muscles, relative to the immediate surrounding non-vocal motor arcopallium (**Supplementary Fig. 1a**). For humans, prior studies showed that *PLXNA1* was downregulated in primary motor cortex (M1) layer 5 neurons^14^, but did not examine the speech laryngeal motor cortex (LMC). A recent study used brain coordinates of microarray expression data from the Allen Human Brain Atlas and reported that *PLXNA1* was downregulated in the dorsal LMC (dLMC) speech region relative to non-speech motor cortex (**Supplementary Fig. 1b**).^11^ Human dLMC has the highest specialized gene expression convergence to songbird RA among vocal learning brain regions. Moreover, *PLXNA1* was significantly downregulated relative to mean cortical expression, in the dorsal laryngeal sensory cortex (dLSC) and overall LMC. Thus, along with the findings of Gu and colleagues (2017)^14^ for forelimb brain circuits, the expression patterns reported here support *PLXNA1* as a strong candidate for testing the direct vocal cortico-motoneuronal hypothesis.

### Mice with conditional knockdown of *PLXNA1* expression have enhanced cortical projections to vocal motor neurons

We crossed floxed *PLXNA1^fl/fl^* mice^16^ with the *RBP4-CRE* line (**Supplementary Fig. 1c**), the latter of which expresses CRE in layer 5 cortical neurons driven by the *RBP4* promoter (**Supplementary Fig. 1d**), which will conditionally knockout *PLXNA1* expression via excision of exon 2. Reduction of mouse *PLXNA1* mRNA expression in M1 layer 5 neurons was demonstrated by *in situ* hybridization in homozygous knockout (KO), *PLXNA1^fl/fl^;RBP4-CRE* (*PLXNA1^KO^;RBP4-Cre*) mice, relative to wildtype (WT), *PLXNA 1^+/+^;RBP4-CRE* (*PLXNA1^WT^;RBP4-CRE*), control animals at post-natal (P) day 12 (**Supplementary Fig. 1e,f**). We injected an anterograde viral tracer, AAV1.hSyn.tdTomato, into M1/LMC of wildtype and conditional knockout adult mice (**Fig. 1b**). Seven days after injection, we examined the density of tdTomato+ axonal fibers projecting from layer 5 cortical neurons to the brainstem. Consistent with many previous findings^7^, wildtype animals had very sparse to no innervation (red, tdTomato+ labelled fibers) of cortical projections to Amb vocal motor neurons (green, Choline Acetyltransferase, ChAT, labelled; **Fig. 1c-e**). However, mice with reduced *PLXNA1* expression in layer 5 neurons had visibly notable enhancement of cortical fibers innervating Amb (**Fig. 1f-i**). To quantify these results, we measured the density of axonal fibers normalized to the area of AAV1.hSyn.tdTomato viral expression in the cortex at the site of injection. We found that heterozygote and homozygous conditional knockouts had a 0.75 and 1.67-log fold increase in axonal density, respectively, compared to wildtype controls (**Fig. 1j**). No significant differences in axonal density were observed in the nearby reticular formation or entire hemifield of the contra-lateral brainstem (**Fig. 1k,l**).

Previous studies have has shown that some direct cortico-motoneuron projections to forelimb motor neurons in the spinal cord exists in mouse pups soon after birth, but that these are entirely pruned by the time the mice become adults and that reduction of *PLXNA1* expression in the cortex inhibits this developmental pruning^14,17^. We wondered if cortical projections to vocal motor neurons in Amb were regulated by a similar mechanism. We again injected AAV1.hSyn.tdTomato into M1/LMC of post-natal day 5 (P5) wildtype, heterozygous, and knockout mice. We waited seven days until P12, and quantified the level of normalized axonal density in the brainstem. Similar to adults, we found that *PLXNA1^KO^;RBP4-CRE* mice showed a significant increased innervation of Amb relative to wildtype (**Supplementary Fig. 2a-h**). Cortical projections to other brainstem regions were unaffected by genotype (**Supplementary Fig. 2i,j**). Developmentally, the difference in cortical innervation density to Amb between wildtype pups versus adults was marginal, while cortical axonal density in Amb increased significantly in heterozygous and knockouts mice adults compared to pups (**Fig. 1m**). These findings show that in contrast to cortical projection to forelimb motor neurons in the spinal cord, direct cortico-motoneuronal projections to vocal motor neurons are very sparse during post-natal development and the effect of *PLXNA1* reduction in layer 5 neurons appears to result in enhancement, rather than inhibition of pruning, of cortical axons innervating Amb motor neurons. While pruning of cortical projections to vocal motor nuclei could occur at an earlier developmental stage than in the limb motor nuclei, we find that cortical axons are largely absent from Amb as early as P6 in wildtype pups (data not shown).

### Conditional knockouts of *PLXNA1* have similar number of layer 5 neurons with connectivity to laryngeal muscles

We investigated whether the increased axonal density of cortical projections to Amb in *PLXNA1^KO^;RBP4-CRE* mice could be attributed to an increase in the quantity of layer 5 neurons with connectivity to vocal motor neurons innervating the laryngeal muscles. Using a previous protocol we established^18^, the retrograde transsynaptic viral tracer pseudorabies virus expressing red fluorescent protein (PRV-RFP) was unilaterally injected into the laryngeal cricothyroid muscles of P7, P12, and adult mice (**Supplementary Fig. 3a**). At approximately 48 and 72 hours post-inoculation in pups and adults, respectively, we quantified the number of retrogradely labeled RFP+ neurons in M1/LMC and other cortical areas with known connectivity to laryngeal muscles. Incubation times were optimized in pups and adults based upon the minimal length of time required to retrogradely labeled RFP+ neurons in the adjacent brainstem reticular formation or solitary track nucleus at each age. We found no significant difference between genotypes in the number layer 5-Amb projecting neurons in the contralateral or ipsilateral M1/LMC (**Supplementary Fig. 3b,c**), nor in the contralateral or ipsilateral insular cortex (**Supplementary Fig. 2d,e)**. However, we did find a notable increase in adults relative to pups in the contralateral M1/LMC and ipsilateral insular cortex (**Supplementary Fig. 3b,e-g**). Given the regional-specificity of the developmental differences, which were not observed in the other two brain regions analyzed, we speculate that a subset of vocal motor circuits undergo an extended period of post-natal maturation. These findings suggest that: 1) the increased cortical innervation to Amb in *PLXNA1^KO^;RBP4-CRE* mice is not due to an increase in the number of layer 5 innervating neurons, but rather, an increase in the axonal density of existing projection neurons; and 2) that cortical connectivity to vocal motor neurons controlling the laryngeal muscles continues to develop into adulthood in wild type animals.

### Adult *PLXNA1^KO^;RBP4-CRE* mice exhibit shorter laryngeal muscle activation latencies after cortical stimulation

The presence of increased direct projections does not necessarily mean that they are functional. If functional, one would expect that stimulation of the layer 5 cortical neurons should result not only laryngeal muscle contractions, but contractions that on average are faster than wildtype. To test function, we utilized a protocol on paired intracortical microstimulation (ICMS) with electromyographic (EMG) recordings from the cricothyroid muscle of the larynx and the extensor carpi radialis in the forelimb^8^. In anesthetized animals, single biphasic stimulation pulses were delivered to layer 5 at sites across the motor cortex, with currents from 50 to 450 μA. Two microwire fishhook electrodes were inserted in each muscle, and EMGs from both muscles were recorded using a differential amplifier (**Fig. 2a**). At each single site and for each individual ICMS current, muscle EMGs were averaged and aligned to the stimulus time, creating a stimulus-triggered average (StTA). We measured the pre-stimulus mean and standard deviation across stimulus events, and set the EMG threshold as 2SD above the pre-stimulus mean. We calculated the response latency as the first time at which the post-stimulus StTA exceeded this threshold for at least 0.5 ms (**Fig. 2b**). For each site at which any EMG(s) occurred, we calculated the latency at the minimum stimulation current which elicited a muscle response. We found, on average, that *PLXNA1^KO^;RBP4-CRE* animals had a 2-fold shorter M1-to-CT latency relative to wildtype, and that this effect appeared to be driven most by the middle and posterior (e.g. LMC) regions of the motor cortex (**Fig. 2d-e**). The reduced latency in the 10-20 ms range is consistent with activating layer 5 neurons that have more monosynaptic connections to Amb, in addition to the maintained indirect connections to the surrounding reticular formation. Surprisingly, in these same animals we did not see an overall decrease in the latency to activate forelimb muscles that had been reported by Gu and colleagues^14^ with *EMX1-CRE* conditional knockouts of *PLXNA1* (**Fig. 2d**). These results suggest that the increased direct connections from layer 5 to Amb in the *PLXNA1^KO^;RBP4-CRE* animals are functional, and that this increase is relatively specific for vocal motor neurons.

**Fig. 2.**
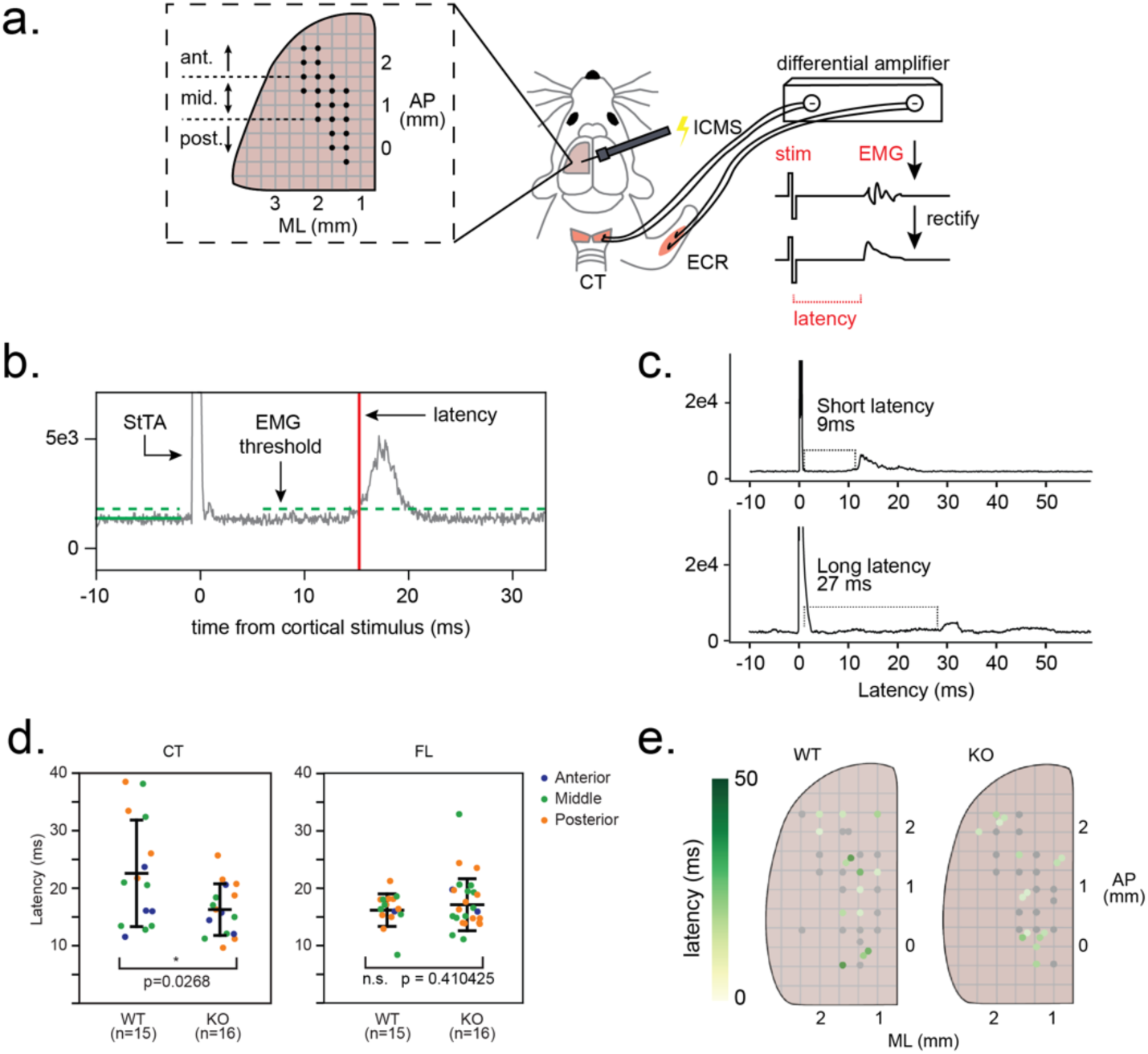
Electrophysiological stimulation of cortico-laryngeal and cortico-forelimb muscle connections in *Plxna1-Cre* adult mice. **a**, Illustration of ICMS-EMG experimental configuration. Left, location of intra-cortical stimulation regions indicated by dotted line inset with coordinates relative to Bregma. Center, ICMS and EMG surgical prep for cricothyroid and extensor carpi radialis muscles. Right, EMG recording and signal processing. **b**, Example cricothyroid EMG stimulus triggered time average (StTA, grey): mean of all ICMS-EMG trials at a single current for a single cortical site, aligned to stimulus. Green solid line: pre-stimulus mean. Green dotted line: EMG threshold (2SD above pre-stimulus mean). Red solid line: latency (first post-stimulus time point at which StTA exceeds EMG threshold). **c**, Example EMG responses with shorter (9ms, above) and longer (27ms, below) latencies. **d**, For each site, latency of StTA at minimum current at which an EMG could be evoked. Left, mean latency for cricothyroid (CT) EMG responses was significantly lower in *PLXNA1^KO^;RBP4-CRE* mice compared to *PLXNA1^WT^;RBP4-CRE* controls (Ind t-test with unequal variance: p=0.02681; t=2.39046; df=19.93167; n=16 sites across 5 KO animals; n=15 sites across 5 WT animals). Right, mean forelimb (FL) EMG latency was not significantly different between *PLXNA1^KO^;RBP4-CRE* and *PLXNA1^WT^;RBP4-CRE* mice (Ind t-test with equal variance: t=-0.74830; p=0.45865; df=40; n=26 sites, N_KO_=5 MICE; n=16 sites, 5 N_WT_=5 mice). Color, site location (anterior, middle, posterior; see panel a). **e**, Spatial distribution of cricothyroid EMG latencies across all stimulation sites for *PLXNA1WT;RBP4-CRE* and in *PLXNA1^KO^;RBP4-CRE* mice with coordinates relative to Bregma. Point color, latency (see color bar, left). Grey, non-responsive site. ‘*’ p<0.05.

### Genotype-dependent changes in vocal plasticity, complexity, and coordination across social contexts

To test whether reduction of *PLXNA1* in layer 5 neurons in *PLXNA1^KO^;RBP4-CRE* mice is associated with any effect on vocal behavior, we analyzed the ultrasonic vocalizations (USVs) of adult male songs, as they are known to show variation in different social context^19^. Following our previous protocols^19^, adult male mice were habituated with an adult live female for 12 hours prior to recording vocal behavior. Each male was randomly assigned one of four stimulus conditions to evoke production of USVs: anesthetized male (AM), anesthetized female (AF), female urine (UR), or live female (LF). USVs were recorded during five minute sessions under each condition for three consecutive days. We then performed analyses on acoustic features of song bouts and categorized individual USVs within song bouts into five syllable types based on frequency jumps (non-jump simple ‘s’, up-jump ‘u’, down-jump ‘d’, multi-jump ‘M’, or unclassified ‘UC’).

We sought to assess the impact of genotype on acoustic features of song bouts at a high-level view using principle component analysis (PCA) applied to all spectral-temporal-informational features extracted from male vocalizations under each social context. We defined song bouts as sequences of five, or more, USV syllables separated by less than 250 ms of inter-syllable silence^19^. We found that the *PLXNA1^KO^;RBP4-CRE* mice had greater variance across song bouts in the AM context, especially along the first principle component (PC1), but less variance in the AF context compared to *PLXNA1^HET^;RBP4-CRE* or *PLXNA1^WT^;RBP4-CRE* controls **(Fig. 3a,b**). Song bouts in the UR and LF context were largely overlapping (**Fig. 3c,d**). Examining the extent to which each acoustic feature loaded onto each principle component, we found that PC1 variance in the AM context was best explained by acoustic features of average bandwidth, velocity, and covariance (CV) of USV syllables per song bout, whereas PC2 was best explained by temporal features of bouts, such as bout duration/length and informational encoding measured by Shannon’s entropy, respectively (**Fig. 3e**). In contrast, PC1 for the AF context had the opposite relationship, or negative correlation, with many of these acoustic variables. Variable loading in the AF context for PC2 had similar rank order and directionality of effects as those in the AM context (**Fig. 3f**). However, when comparing individual acoustic features alone, we find more limited differences across genotypes and social contexts (**Fig 4a-d, Supplementary Fig. 4a**). Only song bout duration and entropy were significantly higher in the AM context for homozygous and heterozygous mutants (**Fig. 4e-i**). These results suggest that the overall genotypic differences in vocal plasticity between AM and AF contexts are produced through the combined effects of acoustic features. In the combined effects, *PLXNA1^KO^;RBP4-CRE* conditional knockouts exhibited more pronounced changes in vocal song bout plasticity across social contexts that involve the presence of an anesthetized conspecific compared to wildtype mice.

**Fig. 3.**
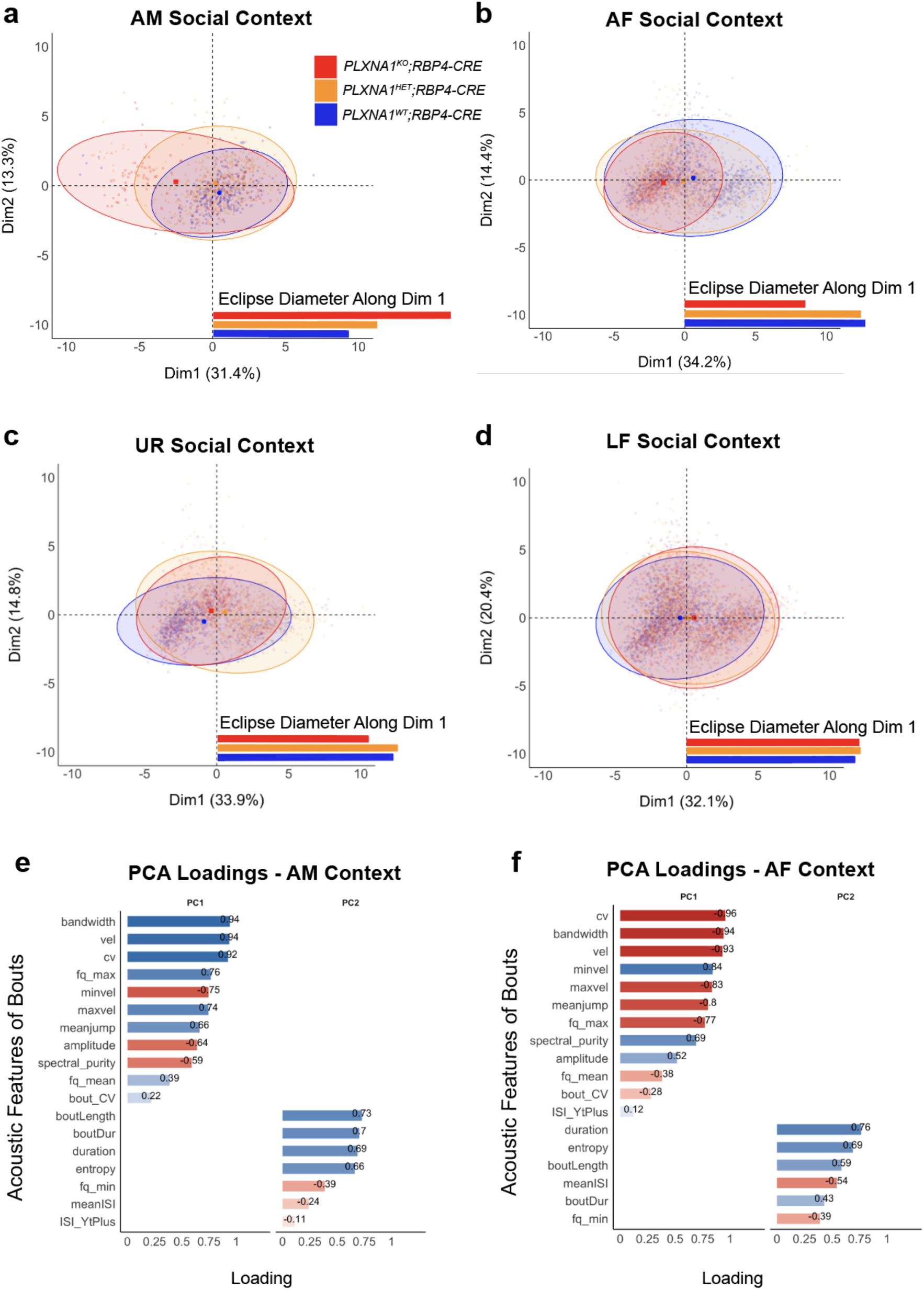
Plasticity of song bouts and acoustic features of *Plxna1-Cre* adult mice across social contexts. **a-d,** Principal component (PC) analysis of 12 acoustic features, six temporal features and Shannon’s entropy performed on song bouts recorded during five minute sessions for conditional knockouts of *PLXNA1* adult mice (N_WT_=7, N_Het_=12, N_Ko_=7) under four different social contexts: anesthetized male (AM), anesthetized female (AF), female urine (UR), or live female (LF). Shaded eclipse areas for *PLXNA1^WT^;RBP4-CRE* (blue), *PLXNA1^HET^;RBP4-CRE* (orange), and *PLXNA1^KO^;RBP4-CRE* (red) are visualized at the 95% confidence level with centroids representing the centered mean for each eclipse colored by genotype. The diameter of each eclipse, along dimension 1 (x-axis), is provided for comparison. **e**,**f**, Loading weights of each acoustic-temporal-informational feature provided for the first two PCs with no rotation in the AM and AF contexts. Colors represent degree of positive (blue) or negative (red) loading or correlation between the variable and contribution to the respective PC.

**Fig. 4.**
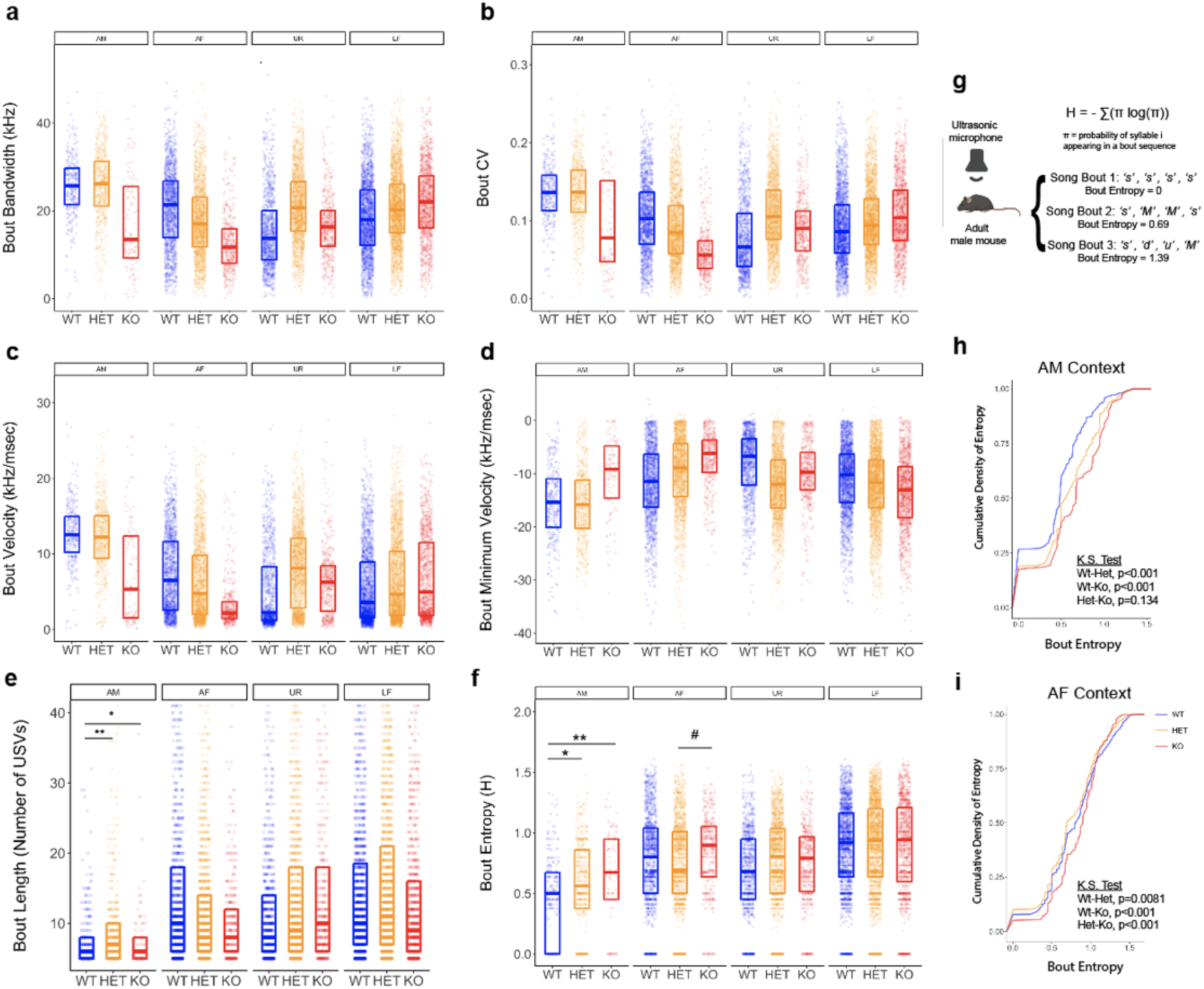
Analysis of individual acoustic features of song bouts in *PLXNA1* conditional knockout adults. **a**-**f**, Post hoc analysis of six individual acoustic features with heavy loadings onto the first two PCs. Boxplots capturing the lower 25^th^ and upper 75 quartiles with median values (colored horizontal line) are provided. The bandwidth, covariance (cv), velocity, and minimum velocity of USVs averaged for each song bout were not significantly influenced by genotype, but bout length was significantly longer in heterozygotes (p=0.00178, df=51.4, t=-3.64) and slightly shorter in homozygous mutants (p=0.0317, df=86.7, t=-2.57) compared to wildtypes. Entropy of bouts (**f**,**g**) vocalized in the AM context was higher in *PLXNA1^KO^;RBP4-CRE* (p=0.0042, df=41.5, t=-3.403,) and *PLXNA1^HET^;RBP4-CRE* (p=0.0106, df=31.2, t=-3.110) mice compared to *PLXNA1^WT^;RBP4-CRE* controls. **h**,**i**, Cumulative density distribution plots for entropy in the AM and AF contexts show wildtype mice accumulate more low entropy bouts in the first quartile while *PLXNA1* conditional knockout mice accumulate significantly more high entropy bouts in the upper quartiles of the distribution. Comparison of cumulative density distributions used two-sided Kolmogorov-Smirnova test for significance. ‘***’ p<0.001, ‘**’ p<0.01, ‘*’ p <0.05, ‘_#_’ p <0.1.

To assess possible relationships or coordination between spectral-acoustic and temporal-informational features of song bouts, we performed a Pearson’s correlational analysis. We observed many expected correlations between features (bandwidth and velocity) that were consistent across groups (**Fig. 5a-c; Supplementary Fig. 5**); but several that were specific to conditional knockout mice, in the AM context. In particular, song bout length and song entropy showed no correlation with spectral purity of song bout syllables in wildtype mice, a weak positive correlation in heterozygotes, and a strong positive correlation in homozygous mice (**Fig. 5d,e**). Conversely, the covariance of syllable ISIs and frequency modulation had the opposite relationship with spectral purity, a strong negative correlation in the homozygous mice (**Fig. 5f,g**). In general, in the AM context, song bouts of *PLXNA1^KO^;RBP4-CRE* mice exhibit strong pairwise correlations that suggest stronger coordination, or coupling, of acoustic features within song bouts compared to wildtype mice. The correlational structure of song bout acoustic features in other social context show different patterns (**Supplementary Fig. 5**).

**Fig. 5.**
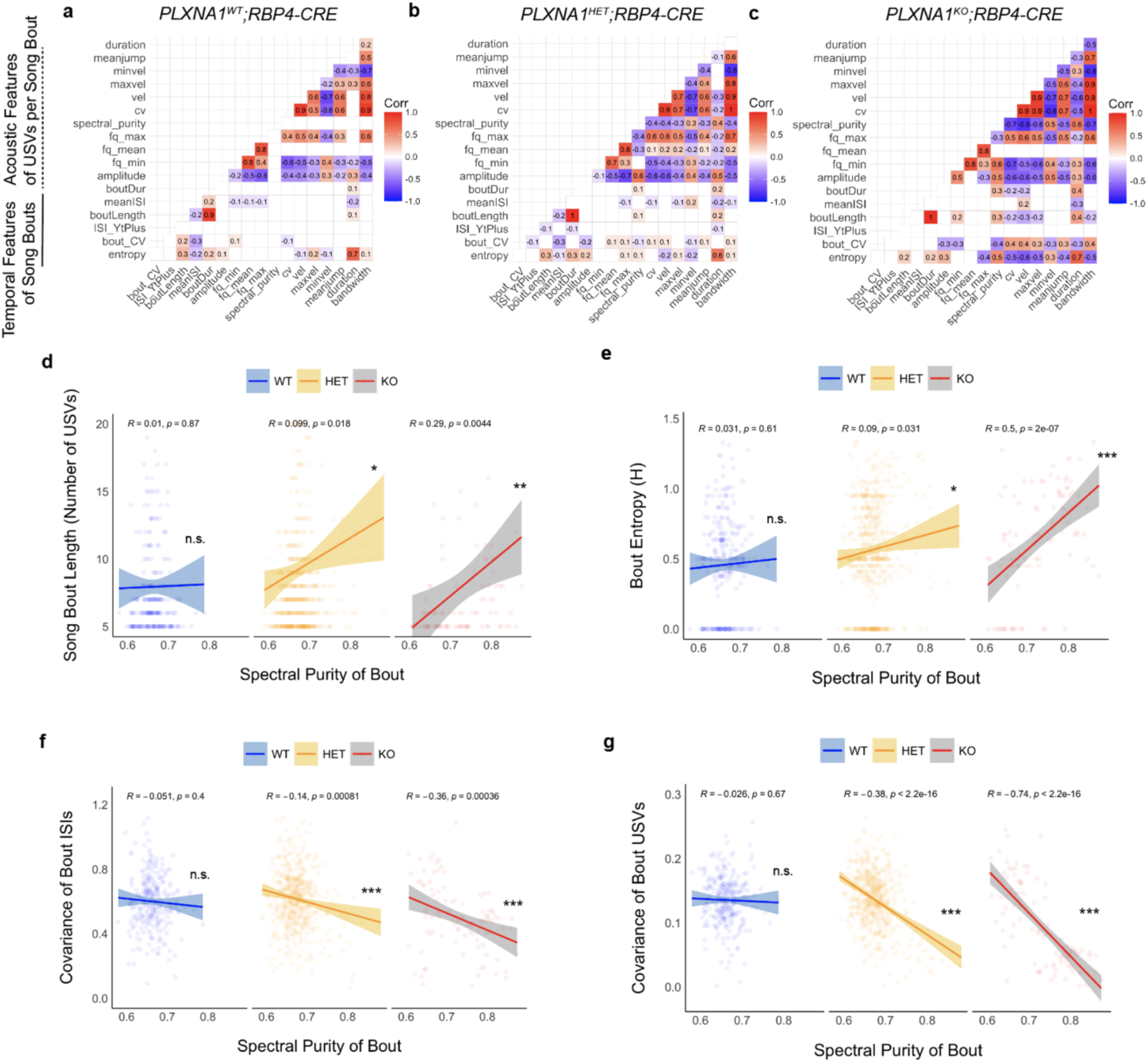
Correlation of acoustic-temporal-informational features of adult song bouts. **a-c**, Correlational structure of song bout features analyzed in adult males (N_WT_=7, N_HET_=12, N_KO_=7) vocalizing in the anesthetized male (AM) context. Correlational matrices were computed using Pearson’s t-distribution product coefficient method to identify significant positive (red) and negative (blue) correlations. Highly similar correlational patterns were obtained using an alternative rank-based Spearman correlational test (data not shown). **d**-**g**, Relationship between (**d**) song bout length, (**e**) the entropy of bout syllable encoding, and (**f**) covariance (CV) of inter-syllable intervals (ISIs), with spectral purity of USVs in song bouts analyzed using linear regressions to assess the impact of genotype on vocal coordination between features. Confidence intervals cover 95% of data color coded by genotype: *PLXNA1^WT^;RBP4-CRE* (blue), *PLXNA1^HET^;RBP4-CRE* (yellow), *PLXNA1^KO^;RBP4-CRE* (red). Correlational coefficients (R value) and statistical significance of the correlations (p) are presented for each regression. ‘***’ p<0.001, ‘**’ p<0.01, ‘*’ p<0.05, ‘_#_’ p<0.1, ‘n.s.’ non-significant.

We also examined acoustic features at the level of individual USVs per syllable type across social contexts. Most acoustic features of USV syllable types per social context were similar across genotypes (**Fig. 6a, Supplementary Tables. 1,2**). However, we found that the ‘d’ syllable type for most measures and the ‘u’ syllable type for a subset of measures were lower in *PLXNA1^KO^;RBP4-CRE* mice, compared to heterozygous and wildtypes, specifically in the AM context. There were no differences in vocal production or in the relative proportion of syllable types emitted from conditional knockout mice across genotypes (**Supplementary Fig. 3b,c**).

**Fig. 6.**
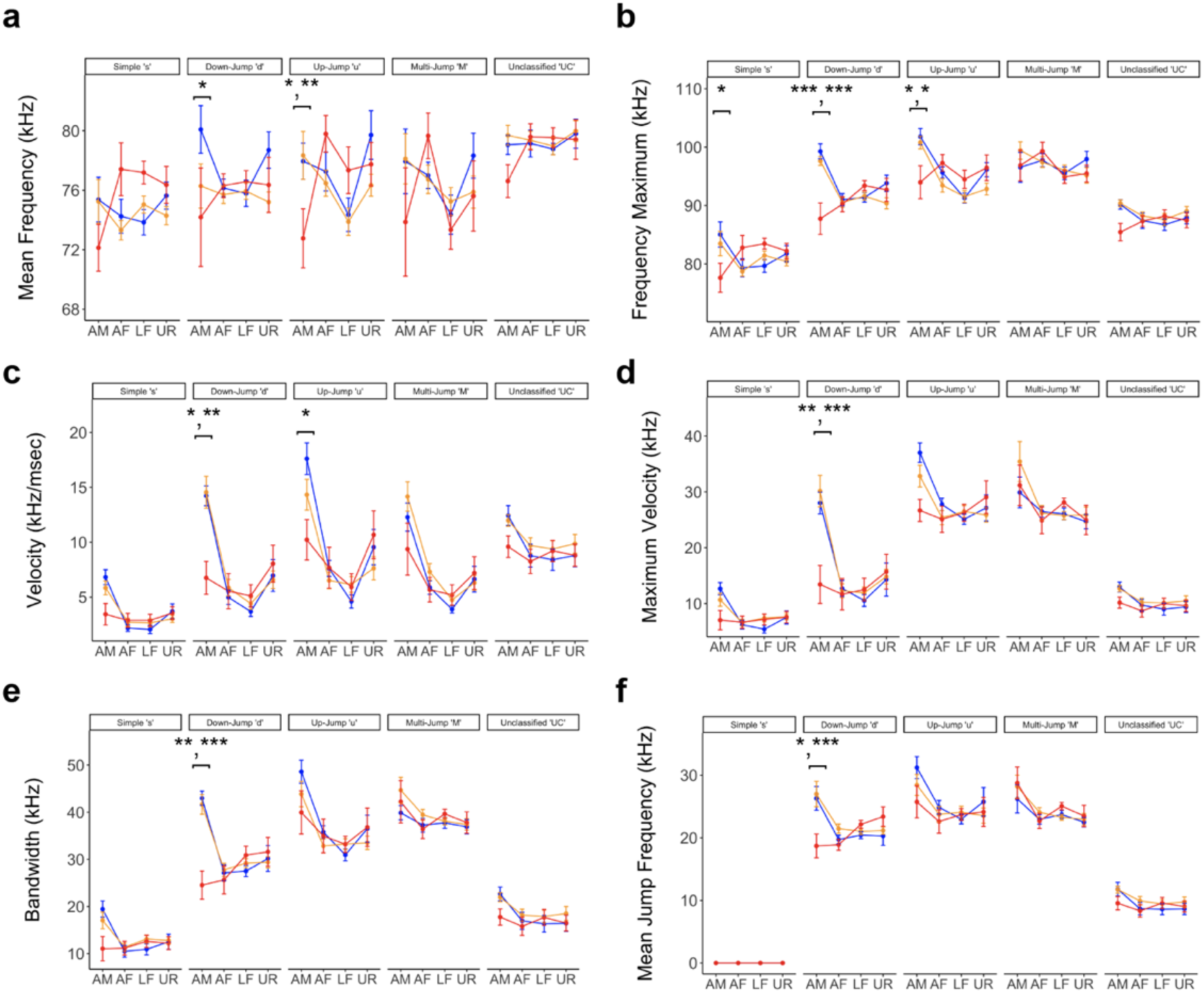
Acoustic features of syllable types detected across social contexts. Analyses of syllable types performed on *PLXNA1* conditional knockout adult male mice (N_WT_=7, N_HET_=12, N_KO_=7) recorded during three five minute sessions in four social contexts. **a,** Mean frequency was significantly different for ‘d’ syllables, WT-KO (p=0.0122, df=477, t=2.86), ‘u’ WT-KO (p=0.0191, df=445, t=2.71) and ‘u’ HET-KO (p=0.0075, df=404, t=3.02) comparisons. **b**, For maximum frequency, significant differences were found for ‘s’ WT-KO (p=0.0187, df=214, t=2.73), ‘d’ WT-KO (p=0.0046, df=279, t=3.83), ‘d’ HET-KO (p=0.00021, df=218, t=4.05), ‘u’ WT-KO (p=0.0307, df=258, t=2.55) and ‘u’ HET-KO (p=0.0231, df=230, t=2.65) comparisons. **c**, Velocity of frequency change for ‘d’ WT-KO (p=0.0417, df=108, t=2.44), ‘d’ HET-KO (p=0.0027, df=88, t=3.42), and ‘u’ WT-KO (p=0.0467, df=101, t=2.41) syllables in the AM context were significant. **d**, Maximum rate of frequency change was significant for ‘d’ WT-KO (p=0.0028, df=206, t=3.34) and ‘d’ HET-KO (p<0001, df=161, t=5.38) in the AM context. **e**, Bandwidth of USVs had significant differences for ‘d’ WT-KO (p=0.00222, df=124, t=3.45) and ‘d’ HET-KO (p=0.000124, df=99, t=4.29). **f**, Significant differences in mean frequency jump included ‘d’ WT-KO (p=0.0220, df=417, t=2.66) and ‘d’ HET-KO (p=0.00027, df=332, t=3.96). Based on spectrogram characteristics, most ‘UC’ syllables are low bandwidth USVs with pitch jumps. Many significant pair-wise differences in acoustic features (13/29 comparisons) were observed with ‘d’ syllables in the AM context (**Supplementary Tables. 1,2**). Linear models included genotype, syllable type, and social context as main effect variables with individual mice as random effects. Post hoc analysis performed with Tukey adjustment for multiple test corrections used to identify significant genotype, social context, and syllable type interactions for each acoustic features. Points represent mean values averaged from each recording session with SEM error bars. ‘***’ p<0.001, ‘**’ p<0.01, ‘*’ p<0.05.

### Mouse pups with conditional reduction of *PLXNA1* exhibit limited differences in isolation calls

We investigated vocal behavior of mouse pups to assess whether the differences seen in vocal behavior of adults with conditional downregulation of *PLXNA1* in layer 5 cortical neurons is present at an earlier age. Because mice pups do not produce courtship songs, we recorded pup vocalizations using a standard maternal-pup isolation paradigm at P7 and P12 in a sound isolation chambers for five minutes. As the bigger differences between knockout and wildtype adults were in acoustic features of song bouts, we assessed whether similar differences could be found in pup isolation call bouts. ‘Isolation bouts’ were defined using similar criteria as those used in adult song bouts: sequences of five, or more, USV syllables separated by less than 250 ms of silence. Using PCA, similar as we did on adults, the centroids and eclipse areas of isolation bouts at P7 were largely overlapping among genotypes (**Fig. 7a**). Song bouts at P12 for conditional knockout mice became slightly differentiated from wildtypes (**Fig. 7b**). The loadings of acoustic features onto each principle component included spectral, temporal, and informational features of isolation bouts (**Fig. 7c,d**). The relative rank of acoustic feature loadings shifted from P7 to P12 with measures of USV frequency modulation within songs bouts (bandwidth, CV, and jump frequency) and informational content (entropy) loading more strongly onto PC1, accounting for 36.9% of bout variance, at the later developmental stage.

**Fig. 7.**
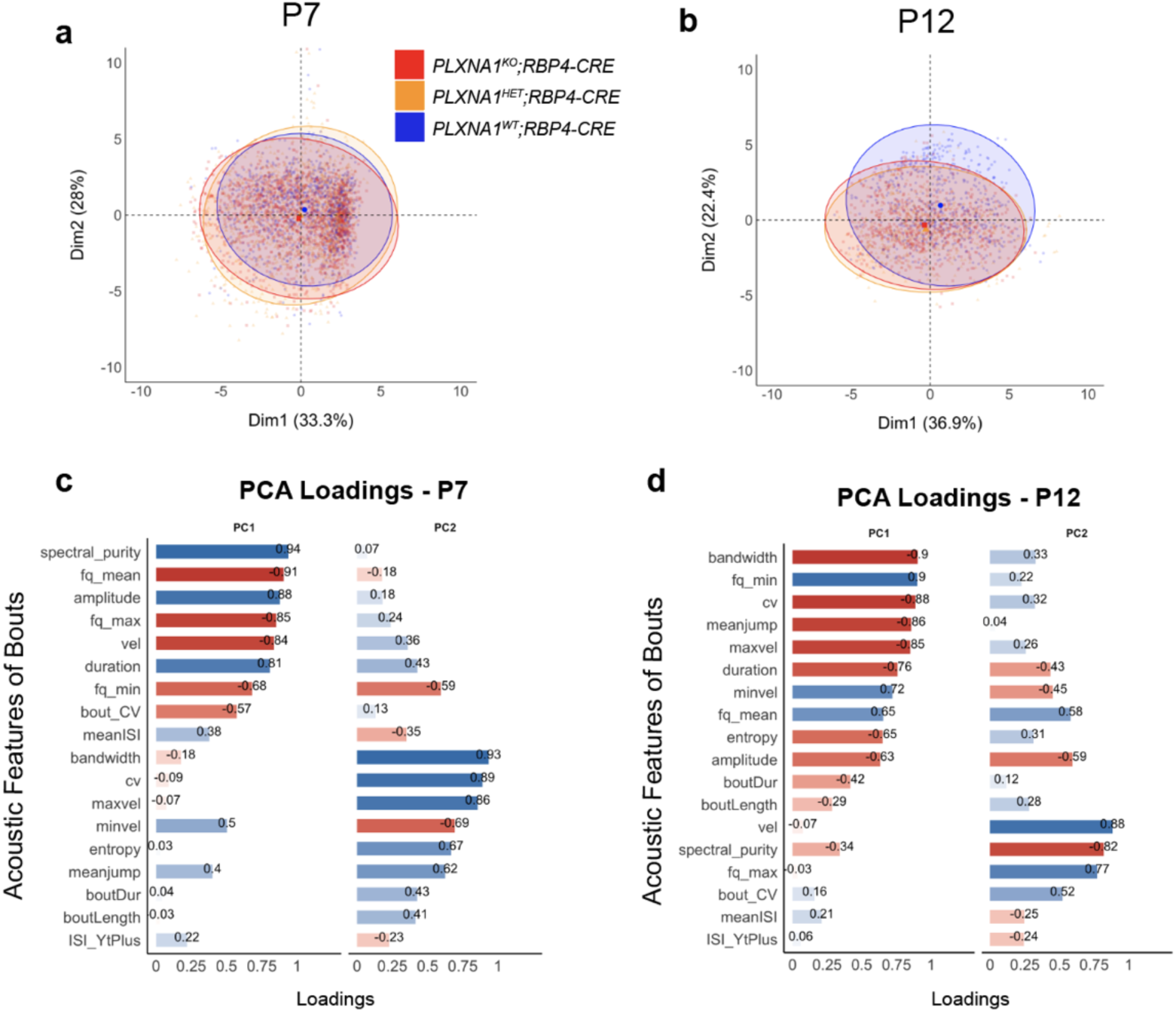
Principal components of acoustic-temporal-informational features of pup isolation bouts. Principal components (PCs) were calculated, similar to adult analysis, using 12 acoustic features and seven temporal features of isolation bouts, and one measure of information encoding (entropy), taken from pup (N_WT_=14, N_Het_=28, N_Ko_=17) USVs recorded during five minutes of maternal-pup separation. **a**,**b**, First two principal components (PCs) visualized for features analyzed from (**a**) P7 and (**b**) P12 pups. Shaded eclipse areas for *PLXNA1^WT^;RBP4-CRE* (blue), *PLXNA1^HET^;RBP4-CRE* (orange), and *PLXNA1^KO^;RBP4-CRE* (red) are visualized at 95% confidence level with centroids representing the mean center for each eclipse. The contribution of each dimension to the overall variance is provided alongside each axis in parenthesis. **c**,**d**, Loading weights of each feature mapped onto PCs are provided with no rotation. Color represents degree of positive (blue) or negative (red) correlation between the variable and principle component.

When studying the acoustic features of individual USVs, we found that vocal production decreased significantly from P7 to P12 (**Supplementary Fig. 4a),** as expected^20^, with no or limited main effects of genotype on the ratio of syllables typed detected or acoustic features (**Supplementary Table. 3**). However, post hoc analysis identified significant pairwise differences localized to ‘u’ syllables emitted by P12 pups (**Supplementary Table. 4**). In particular, spectral purity, amplitude, and duration of USVs emitted from heterozygous conditional knockout pups were significantly higher or longer compared to homozygous animals or wildtype controls. Surprisingly, we observed that mean frequency, minimum frequency, maximum frequency, and velocity had differences only between heterozygous and knockout animals.

Closer investigation of individual isolation bout temporal features also showed strong developmental changes in bout production, length of bouts, mean ISIs of USVs within bouts, but no or limited genotypic effects (**Fig. 8a-c**). However, post hoc analysis revealed that the covariance of ISIs within bouts, latency to initiate bouts, and syllable entropy exhibited strong developmental and genotype-specific effects with conditional knockout pups showing more pronounced trends toward stereotypy and reduced variation (**Fig. 8d-f**). At P7, genotype only had a marginal influence on bout entropy between knockouts and wildtype pups, which became more pronounced at P12. For acoustic features of song bouts, genotype more significantly influenced bandwidth, minimum frequency, covariance of frequency modulation (USV-CV), and mean jump frequency (for syllables with pitch jumps) (**Fig. 9a-d,g-i**), with no apparent influence on spectral purity of velocity of USVs within bouts (**Fig. 9e-f, g-i**). Overall, these findings indicate that relative to wildtype, pups with condition depletion of *PLXNA1* in layer 5 neurons have developmentally emergent differences in song bout features that become more pronounced in the social context vocal behavior of adult animals.

**Fig. 8.**
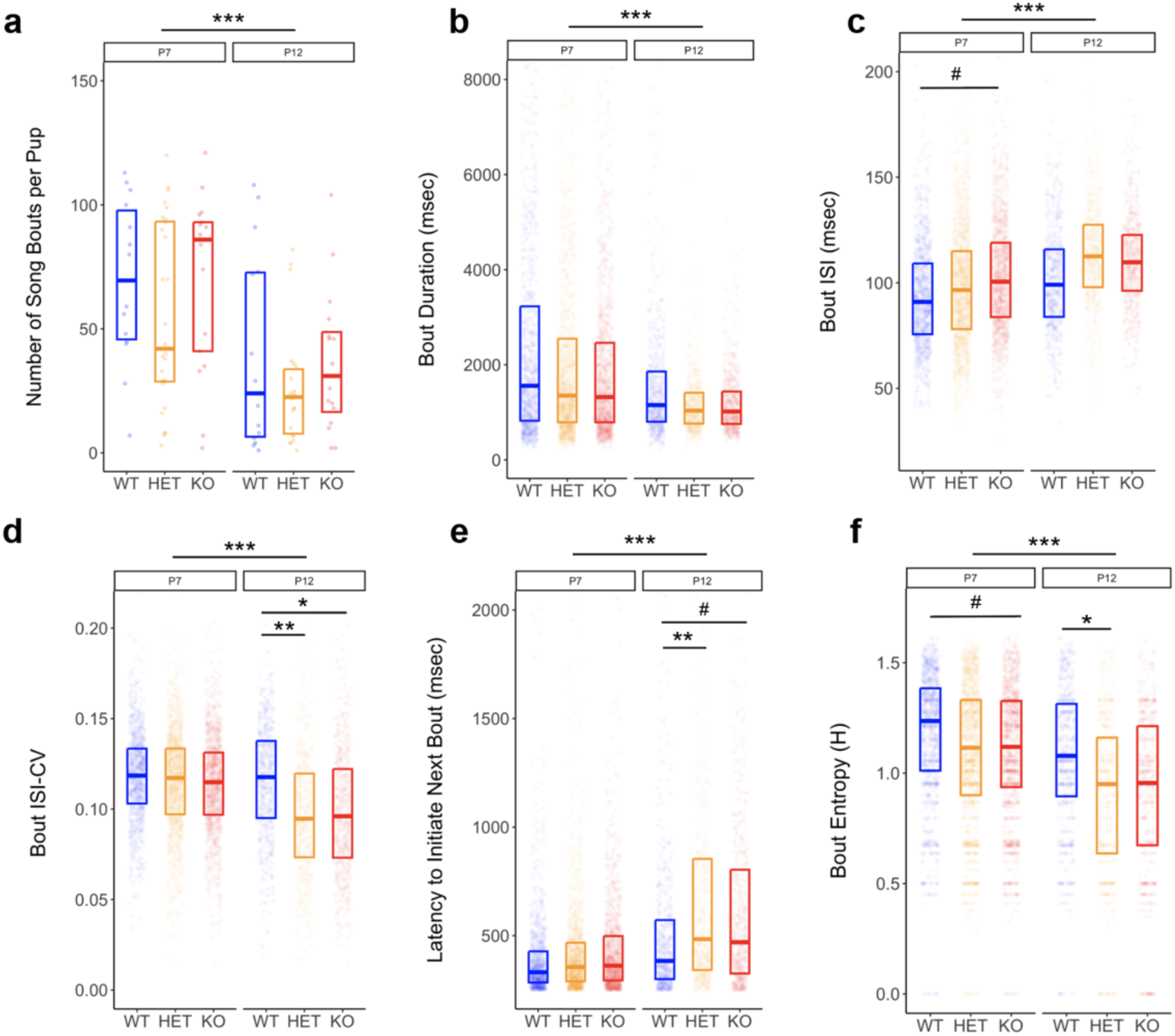
Developmental trajectory of temporal and informational features of isolation bouts. *PLXNA1* conditional knockout mouse pups (N_WT_=14, N_HET_=28, N_KO_=17) were recorded during maternal-pup isolation for five minutes and analyzed at the level of bouts longer than 4 sequences of USVs separated by less than 250 ms silence. **a**, Number of bouts decreased significantly across development (p=8.59×10-6, F=21.9, df=1) with no major effect due to genotype (p=0.120, F=2.16, df=2). **b**, Length of isolation bouts significantly decreased with age (p<2.2×10-16, F=273, df=1). **c**, The mean inter-syllable interval (ISI) of USVs within bouts increased from P7 to P12 (p<2.2×10-16, F=234, df=1) with *PLXNA1^KO^;RBP4-CRE* pups showing marginally higher mean ISIs at earlier stages (p=0.0639, t.ratio=-2.30, df=56.8). **d**, Covariance (CV) of ISIs within bouts decreased in P12 pups compared to P7 (p<2.2×10-16, F=212, df=1) with genotype having a marginal main effect (p=0.0852, F=2.58, df=2). Post-hoc comparisons revealed that *PLXNA1^KO^;RBP4-CRE* (p=0.0248, t.ratio=2.68, df=67) and *PLXNA1^HET^;RBP4-CRE* (p=0.0069, t.ratio=3.137, df=73) mice had lower variation in ISIs at P12 compared to *PLXNA1^WT^;RBP4-CRE* pups. **e**, The amount of time between bouts, or latency to initiative a bout, increased from P7 to P12 (p<2.2×10-16, F=418, df=1) with a marginal main effect of genotype (p=0.0683, F=2.83, df=2). Latencies in wildtype P12 pups were significantly lower than *PLXNA1^HET^;RBP4-CRE* (p=0.0055, t.ratio=-3.202, df=78.1) and *PLXNA1^KO^;RBP4-CRE* (p=0.10, t.ratio=-2.064, df=70.3) pups. **f**, The entropy of isolation bouts decreased in older pups (p<2.2×10-16, F=317, df=1). Bout entropy was marginally lower in *PLXNA1^KO^;RBP4-CRE* (p=0.07, t.ratio=2.241, df=55.6) compared to wildtype pups at P7, while *PLXNA1^HET^;RBP4-CRE* entropy was significantly lower at P12 (p=0.0429, t.ratio=2.472, df=56.5). Significance assessed with linear mixed models using age and genotype as main effect variables and random effects of individual pups. Tukey adjustment for multiple tests used in post hoc analysis. ‘***’ p<0.001, ‘**’ p<0.01, ‘*’ p <0.05, ‘_#_’ p<0.1.

**Fig. 9.**
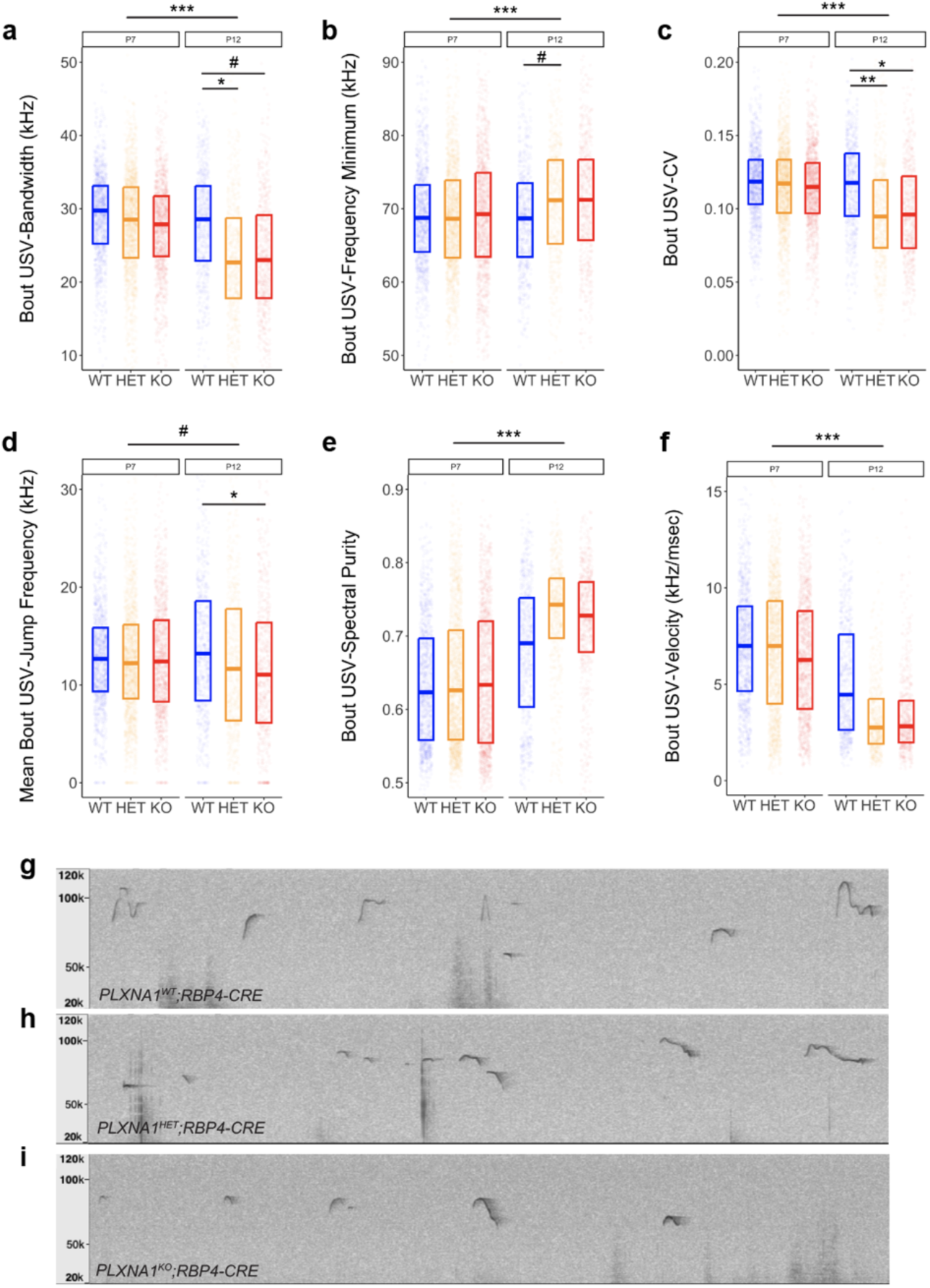
Acoustic features of isolation bouts in conditional pup knockouts. Vocal behavior of *PLXNA1* conditional knockout pups (N_WT_=14, N_HET_=28, N_KO_=17) were recorded and analyzed as described in the methods. **a**, The bandwidth of isolation bouts was significantly lower in *PLXNA1^KO^;RBP4-CRE* (p=0.0732, t=2.23, df=63.4) and *PLXNA1^HET^;RBP4-CRE* (p=0.0137, t=2.90, df=67.4) mutants compared to *PLXNA1^WT^;RBP4-CRE* controls. **b**, Marginal increases in the minimum frequency of USVs within bouts of *PLXNA1^HET^;RBP4-CRE* (p=0.0578, t=-2.32, df=84.5) pups were found. **c**, The covariance (CV) of bout USVs was significantly lower in *PLXNA1^KO^;RBP4-CRE* (p=0.0248, t=2.68, df=67) and *PLXNA1^HET^;RBP4-CRE* (p=0.0069, t=3.14, df=73) mutants compared to *PLXNA1^WT^;RBP4-CRE* controls. **d**, Mean jump frequency of bout USVs also decreased in *PLXNA1^KO^;RBP4-CRE* (p=0.0293, t=2.60, df=78.9) mice compared to controls. **e**,**f**, Genotype did not influence spectral purity or velocity of USVs in isolation bouts. **g**-**i**, Representative spectrograms of isolation bouts emitted by each genotype are presented. Significance assessed with linear mixed models using age and genotype as main effect variables and random effects of individual pups. Tukey adjustment for multiple tests used in post hoc analysis. ‘***’ p<0.001, ‘**’ p<0.01, ‘*’ p <0.05, ‘_#_’ p<0.1.

## Discussion

Our study provides the first known case of experimentally inducing the formation of a specialized connection found in vocal learning species in an otherwise vocal non-learning species. Our key findings are: that experimental down-regulation of the axon guidance receptor *PLXNA1* in motor cortex layer 5 neurons, partly recapitulating the specialized pattern in humans and songbirds, results in increased direct innervation to vocal motor neurons, supporting the repulsion hypothesis for the formation of this connection^10^; the newly formed direct projections are associated with decreased latency from the cortex to activate laryngeal muscles; and increased complexity of vocal behaviors (context-dependent plasticity, coordination, informational encoding) in different social context, in support of the Kuypers-Jurgens hypothesis^21^; and these vocal complexity changes occur gradually over post-natal development. These findings provide insight into the molecular and developmental mechanisms that regulate formation of vocal cortico-motoneuronal circuits and an opportunity to examine the biological and behavioral impact of acquiring this connection in a vocal non-learning animal.

As a component of the forebrain vocal learning pathway, the motor cortex-to-brainstem vocal motor neuron circuit has long been argued to play a critical role in the evolution of vocal learning by enabling voluntary control over vocal muscle movements^22,2,4,23,3^. However, the relevance of this neural circuit to vocal learning has been contested^24^, in part, due to the inability to assess the impact of acquiring a new direct motor control circuit on vocal behavior, complexity, plasticity, and other evolutionarily derived subfeatures of vocal learning, in a background of an ancestral vocal non-learning brain. Our study is a first step to experimentally assess this connection’s impact on vocal learning behavior in the context of a vocal non-learning species.

Our attempt to modulate this connection by manipulating the expression of *PLXNA1* was inspired by the specialized downregulation of this candidate gene in the direct projection neurons that control vocalizations in songbirds and humans^9,11,25^, and the successful enhancement of cortico-motoneuronal circuits regulating forelimb dexterity in mice^14^. The latter study found that EMX1-mediate knockdown of *PLXNA1* in the cortex acts to inhibit stereotyped pruning of cortical axons to forelimb motor neurons, transiently formed during development in mice^17^. We restricted the down regulation to layer 5 neurons and instead of inhibition of pruning, we found a gain-of-function phenotype marked by an increase in density of descending cortical axons into the vicinity of ChAT+ motor neurons located in Amb. Moreover, we did not find evidence of increased axonal density in the adjacent reticular formation or overall brainstem hemifield region contralateral to the side of tracer injection, which suggests that loss of *PLNXA1* does not lead to broad formation of supernumerary axons in the brainstem^26^. Thus, we argue that *PLXNA1* acts through two developmental mechanisms to induce the formation of cortico-motoneuronal circuits.

Our findings support the hypothesis that, for vocal learning species, reduced levels of the *PLXNA1* receptor, likely located on the terminals of descending axons^27^, inhibits the responsiveness of descending pallial axons to repulsive ligands expressed by phonatory motor neurons in the brainstem (**Fig. 1a**). The repulsive cues would be expected to be either semaphorin (SEMA) or SLIT ligands^27^. Differential expression of SEMA and SLIT ligands and other SLIT receptors (ROBO) have been found in vocal learning brain regions, as well as high levels of SLIT3 in the Amb equivalent of songbirds, nxIIts^9,27^. It is likely that additional manipulation of these and other axon guidance genes to recapitulate the songbird and human expression patterns, could enhance this connection further or enhanced/induce other specialized connections more so, and lead to more advanced vocal changes.

Vocal learning has historically been viewed as dichotomous trait where species were classified as vocal learners or vocal non-learners. Yet there is a growing recognition that the degree of vocal learning is sufficiently varied across species to justify a less categorical schema, that is, the continuum hypothesis of vocal learning^12,13^. Based on this more permissive view, others have argued for a more modular or multi-dimensional approach when assessing the capacity for vocal learning across species^24,28^. Our findings that an 0.7-1.7-fold increase in the cortico-motoneuron innervation of Amb leads to some changes in vocal behavior complexity with subfeatures found in wildtype mice is consistent with the continuum hypothesis. Moreover, the social context-dependence of these features demonstrate the kind of vocal production variability, or the ability to change the relationship or variability of acoustic features, that are argued to be a subfeature of vocal learning that emerged in more naturalistic settings from what Martins and Boecke would call ‘functional pressures’^24^.

Limitations of our study to consider are that although *RBP4-CRE* expression is highly selective to layer 5 cortical neurons within the forebrain, like all reporter CRE drivers, there are some other cell types the reporter is expressed in^29^. The vocal behavioral changes we observed could be possibly due to changes of *PLXNA1* in other cell types outside of forebrain layer 5 neurons. However, of the cell types known, and the expression of *PLXNA1*, we do not know of any other brain cell type or brain area that expresses RBP4 and overlaps with *PLXNA1* expression. Another consideration is that we downregulated *PLXNA1* in layer 5 neurons throughout the cortex, not just in the LMC region. In songbirds and humans, *PLXNA1* downregulation does occur throughout the equivalent non-vocal layer 5 neurons, but is further downregulated to a greater level in the vocal regions. Lastly, *RBP4-CRE* layer 5 neurons project to a number areas outside of the Amb^30^. Future studies could develop methods to downregulate *PLXNA1* only in neurons that project to Amb and the surrounding reticular formation. At the behavioral level, although we found changes in vocal complexity and in the direction expected for some features, we did not test for vocal imitation abilities. This is difficult to do, considering mice do not have advanced vocal imitation abilities and there is no behavioral paradigm with which to test them. We are working on developing such a paradigm. Regardless of the potential future outcome of those results, the vocal behavioral findings of this study stand alone in demonstrating a phenotype not found in wildtype mice. The impact of acquiring the forebrain-to-phonatory muscle connection to vocal learning, whether viewed along a continuum, contiguum, or modules, can now be examined experimentally in the laboratory.

## Material and Methods

### Mouse lines

Animals were treated according to the ethical standards defined by the National Institutes of Health guidelines and The Rockefeller University (RU) Institutional Animal Care and Use Committee (IACUC) for animal health and care in strict compliance with all recommendations. All efforts were made to minimize animal discomfort and to reduce the number of animals used. Stock C57BL6/J and B6.Cg-*Gt(ROSA)26Sor^tm14(CAG–tdTomato)Hze^*/J (Ai14) mice were obtained from the Jackson Laboratory. Adult *PLXNA1^fl/fl^* mice were obtained from Dr. Yoshida Yutaka^16^, which contain loxP sites 5’ upstream of exon 2 and 3’ downstream of exon 2, and crossed to the *RBP4-CRE* line maintained at RU to generate *PLXNA1^fl/fl^;RBP4-CRE* transgenics (*PLXNA1^KO^;RBP4-CRE*). Transgenic mice were validated by PCR genotyping (primer pairs: 5’-TGGTGCATCGAAGCAACTGGCACTC-3’ and 5’-CACGGCCTCCTCATTCTCTGAGCTA-3’ of the *PLXNA1* locus (and in-situ hybridization of the *PlxnA1* messenger RNA in brain sections. *PLXNA 1^fl/fl^* stock was maintained on C57BL6/J background. Room ventilation, temperature and humidity were controlled with a 12/12 light-dark cycle.

### Viral injections

Adult (P60-P100) and neonate (P0-P5) mice were unilaterally injected with AAV1.Syn.TdTomato in primary motor cortex, using a pulled glass pipette with Monoject II pump at 1nL/sec. Adults were anesthetized using isoflurane, placed in a stereotaxic frame and the skull was exposed to received two 100 nL injections, one at 0.5 AP, +/- 1.4ML, −0.8 DV, and one at 0 AP, +/- 1.4ML, −0.8 DV, coordinates of the M1/LMC region confirmed through pilot experiments with retrogradely transsynaptic tracer PRV-RFP injection in laryngeal muscles. Incisions were closed using vetbond, and mice were treated with bupivacaine and meloxicam post-surgery. Neonates underwent cryoanesthesia, and were placed in a custom 3D printed surgery stage in which the body maintained anesthesia in a cold water bath while the head was held steady for injections. Pups received a single 100nL injection through the skin relative to lambda at 2.2 AP, +/- 0.6 ML, -0.4 DV. Pups were placed on a heating pad to recover before being returned to their dam.

### Antibodies and immunostaining

Mice were anesthetized using isoflurane and perfused using ice-cold PBS and 4% PFA, and post-fixed for two days in 4% PFA. In the neonate group, perfusions took place at P12 directly after isolation call recordings. Adults were perfused 14 days post-surgery. Brains were cryoprotected in PBS with 30% sucrose before being flash-frozen in OCT using a dry ice 2-methylbutane bath. Brains were sectioned at 50uM using a Leica cryostat, and stored free-floating in PBS. Injection sites were mounted onto slides to be imaged for endogenous fluorescence, and brainstem sections containing nucleus ambiguus were collected for IHC using the following protocol. At room temperature, sections were post-fixed for 20 mins in 4% PFA, washed 2 x 15 mins in wash buffer, (1X PBS, 1% Tween-20) and blocked for 2 hours in 1X PBS, 1% BSA, and 0.3% Triton-X. Sections were incubated overnight with anti-RFP antibody (Rockland, 600-401-379, 1:1000) and anti-ChAT antibody (AB144P, 1:100, Chemicon) in an incubation buffer (2% BSA, 0.1% Triton-X in 1X PBS). Sections were washed 3 x 15 mins in a wash buffer, and then covered from light and incubated in secondary antibodies (Donkey anti-Rabbit 568 1:500; Donkey anti-Goat Alexa 488; in 1 x PBS, 2% BSA, 1% Tween-20). Sections were washed 3 x 15 mins in wash buffer before being mounted for imaging. All sections were mounted in a Vectashield softset antifade mounting medium with DAPI and sealed with nail polish.

### Imaging and Quantitative Image Analysis

Fluorescence microscopy was done using an Olympus BX61 TRF microscope. Injection sites were imaged at 4X, and brainstem sections were imaged at 10X. All images were quantified using FIJI (ImageJ) software. Injection site size was quantified using FIJI’s area measurement, thresholded to injection site size. To estimate axonal density, regions of interest including the nucleus ambiguus (Amb), reticular formation (RF), and brain stem hemifield, were drawn using DAPI and ChAT staining to demarcate relevant areas for analysis. Data was collected from 3-5 50µm sections per animal. Images were auto-local thresholded using Phansalkar method^31^, and integrated density measurements were collected from regions of interest. To account for differences in viral spread, integrated density measurements were divided by the estimate size of the injection area using 3-5 non-consecutive sections to determine the normalized axonal density measurement. Normalized axonal density was log transformed for subsequent analysis.

### ICMS-EMG recordings

Following the protocol of Vargas et al 2024^8^, **a**nesthesia was initially induced using brief exposure to isoflurane, followed by an IP cocktail of ketamine (100 mg/kg) and xylazine (20 mg/kg), then maintained with a quarter dose of ketamine-only cocktail every 30 minutes. Custom electrodes were used to record EMG signals. Formvar insulated nichrome wire (0.0015” diameter, A-M Systems) was de-insulated on both ends and threaded through a pulled and beveled glass pipette (Drummond Nanojects). A small ‘fishhook’ was made at the end of the wire just past the bevel point; which was used to embed the fishhook into the muscle. Pairs of electrodes were inserted 1-2 millimeters apart in each muscle, parallel to the direction of the muscle fibers. Vetbond Tissue Adhesive (3M) was applied to the insertion site to secure the wire within the muscle. To expose the laryngeal muscles, a midline opening was made between the sternum and chin with the mouse in the supine position. Skin and fat were retracted to expose the sternohyoid muscle, which was separated along the midline, and the right strap was cut or secured with retractors. The membrane over the right cricothyroid (CT) was gently removed, and 2 fishhook electrodes were placed into the muscle and secured with a small amount of Vetbond. Signal quality was confirmed by the presence of a breath-associated EMG. The skin opening was closed, and the mouse was flipped into the prone position. A section of skin was cut along the right forelimb between the wrist and elbow, membrane was removed over the extensor carpi radialis (ECR), and a pair of fishhook electrodes were inserted and secured with Vetbond. EMG recordings were performed with a differential amplifier (Model 1800, A-M Systems) through a system notch and bandpass (10Hz-5kHz) filter. Amplifier output was captured at 20kHz via a recording DAQ (USB-230, Diligent) connecting to a laptop with DAQami (Measurement Computing) software. Signal output was also linked to an oscilloscope and audio monitor (Model 3300, A-M Systems).

After all electrodes were inserted, the mouse was placed in nose and ear bars. Skin was removed over most of the skull and a large craniotomy was performed over left motor cortex using a dental drill, extending (relative to Bregma) 2-3mm anterior, 1mm posterior, and 3mm lateral (see Fig. 2a). Mediolateral (ML) and anteroposterior (AP) stimulation coordinates were measured relative to Bregma. Stimulation sites were arranged in a grid diagonally along motor cortex between 1mm and 2.3mm ML, and between 0mm and 2.6mm AP, separated by 300µm in both ML and AP directions. 0.1mm deviations were sometimes made to avoid puncturing large blood vessels. Sites were stimulated non-sequentially. A 50µm diameter bipolar cluster stimulating electrode (FHC) was lowered 800-900µm ventral to the brain surface, to the depth of Layer 5b in motor cortex. Stimulations were performed using an isolated pulse stimulator (Model 2100, A-M Systems) with currents from 50µA to 450µA, in 50µA intervals. Single pulses were delivered as a 0.2ms biphasic pulse at 2Hz, continuously for 30s, resulting in 60 single-pulse stimulations per current/site. Once data was collected, signals from each recording were filtered with a 5^th^-order 500 Hz high pass Butterworth forward filter, using the sosfilt function in the *scipy* python library. Signals were then full-wave rectified by taking the absolute value of the voltage. For each round of stimulations at a particular site and current, EMG recordings were aligned to each stimulus event, and the mean signal was computed, creating a single stimulus-triggered average (StTA) for each site/current. All recordings were manually inspected to eliminate sessions with obvious artifacts, including electrical noise or breathing EMGs (in the case of the cricothyroid). To determine response latencies, we first computed the mean and standard deviation of the StTA in the 50ms preceding the stimulus artifact. The EMG threshold was set at 2SD above the pre-stimulus mean. In the post-stimulus period, after a buffer window of 5-7ms (to avoid counting stimulus artifact extended by the forward filter), if the StTA exceeded the EMG threshold for >0.5ms, we counted it as an EMG contraction. The first time-point to cross this threshold was the response latency for the particular site and current. Because at a given site, multiple currents could induce an EMG response, we took the latency from the EMG response at the lowest possible current. All analysis was performed with Python, and with open-source libraries developed for Python, including numpy, pandas, seaborn, SciPy, and matplotlib.

### Recording and extraction of mouse ultrasonic vocalizations

Adult mice were maintained in home cages with 1-3 other male mice prior to vocal recording. Pups were kept with the dam till weaning age at P21. To assess adult vocal behavior, recordings were performed in sound isolation chambers as previously described (see^32^ for details). Briefly, ultrasound microphones were suspended 10 cm above the recording chamber floor. Vocalizations were recorded using Avisoft Bioacoustics equipment and software (Berlin, Germany), including Avisoft UltraSoundGate 416H 1.1 hardware and Avisoft Recorder USG software with a 250 kHz sampling rate, 15 kHz high-pass filter, 0.032 sec buffer, and 256 FFT with 16 bit format for five minutes. Spectrograms were analyzed with a python implementation of Mouse Song Analyzer^33^ (MSA) with the following parameters: purity bandwidth (1000 Hz), snr threshold (10dB), purity threshold (0.5, discontinuity threshold (2), frequency threshold (40 kHz high pass), merge threshold (15 ms), within the frequency range of 110-35 kHz)^7,32,33^. Shannon’s entropy was calculated with a base 1 logarithm in R (*DescTools*) using a table of syllable types (‘s’, ‘d’, ‘u’, ‘M’, ‘UC’) vocalized by an animal during a recording session as the expected amount of information, or number of bits required to encoding the sequence of syllables^34^.

### Statistical analyses

Numerical analysis was conducted in RStudio (Version 1.1.463) with R (4.0.3). Cluster analyses conducted with R packages *FactorMineR*, *factoextra*, *parameters* on default settings with scaled units for all acoustic features. The first two principal components were used for downstream analysis with maximum loading threshold and no rotation. The mean value acoustic features were calculated per animal per recording session and analyzed as response variables in linear mixed models (*lme4*) with genotype, age or social context, and syllable type as main effect predictors. Individual animals with more than one data point (for example, the same animal recorded at P7 and P12, or across social contexts) were modeled as random effects with repeated measures. Main effects were assessed with Type III Analysis of Variance (ANOVAs) tables using the Satterthwaite’s method. Post hoc comparisons were computed with *emmeans* and Tukey adjustment for multiple test correction.

## Acknowledgments

We thank Cesar Vargas for help with the EMG stimulation study; Dominik Biezonski for preparing an earlier draft of Figure 1a hypothesis and help with writing grant proposals for the project; Yukata Yoshida for generously providing the *PLXNA1^fl/fl^* mice; and the Jarvis Lab neuroengineering team for useful discussions. This work was supported by funds from a NIH Directors Transformative R01 award (R01 OD028000-01), Keck Foundation award, and Howard Hughes Medical Institute (HHMI) funds to E.D.J.

## Author Contributions

J.L.B. and E.D.J. conceived the study and wrote the manuscript. J.L.B., L.K., E.L., and V.Y. performed experiments and analyzed data.

## Competing Interest Statement

The authors disclose no competing interests.

## Supplementary Figures

**Supplementary Fig. 1.**
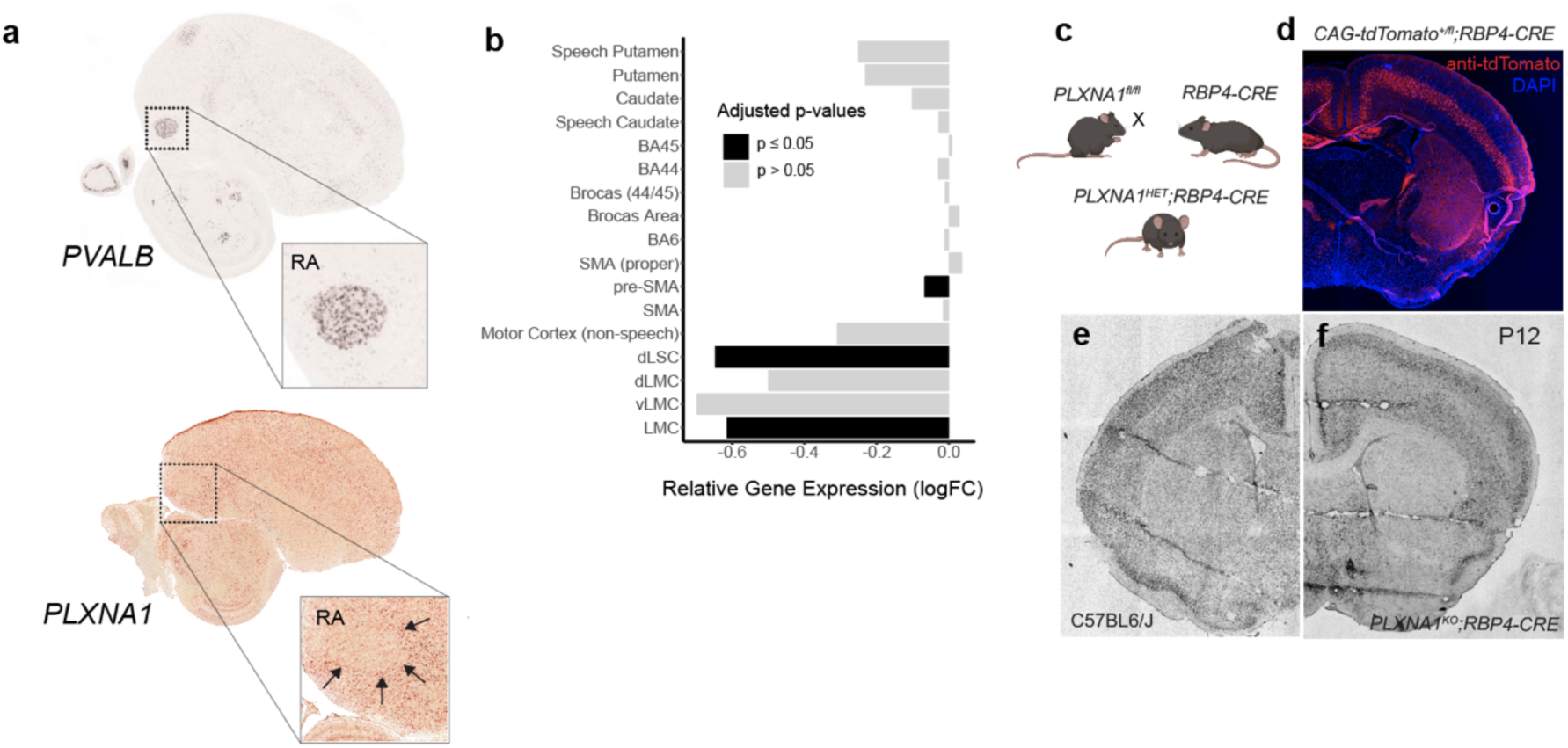
Expression of *PLXNA1* in vocal learning brain regions in humans and songbirds. **a**, Top, location of the robust nucleus of the arcopallium (RA) identified using *in situ* hybridization for *PVALB* obtained from ZEBrA (zebrafinchatlas.org). Bottom, *in situ* hybridization for *PLXNA1* in Zebra finches confirms^11,25^ differential down-regulation in RA (black arrows) relative to the surrounding arcopallium. **b**, Visualization of statistical results from Gedman et. al. (2024) describing genome-wide microarray expression^11^ from the Allen Human Brain Atlas, which show significant (black bars) down-regulation of *PLXNA1* in speech and language-associated areas, including the pre-supplementary area (pre-SMA), laryngeal somatosensory cortex (dLSC) and laryngeal motor cortex (LMC), compared to other regions (gray bars). **c**, Genetic strategy for reducing expression of mouse *PLXNA1* in layer 5 cortical neurons using *PLXNA1^fl/fl^* line crossed to *RBP4-CRE* mice, in order to model down-regulation of human *PLXNA1* in layer 5 neurons of the cortex. **d**, Immunohistochemistry of tdTomato (red) expression activated by a Cre-dependent floxed reporter in P14 juvenile mice. **e**,**f**, *In situ* hybridization for mouse *PLXNA1* mRNA expression in C57BL6J control brains and *PLXNA1^KO^;RBP4-CRE* (*PLXNA1^fl/fl^;Rbp4-Cre*) mutant mice at P12.

**Supplementary Fig. 2.**
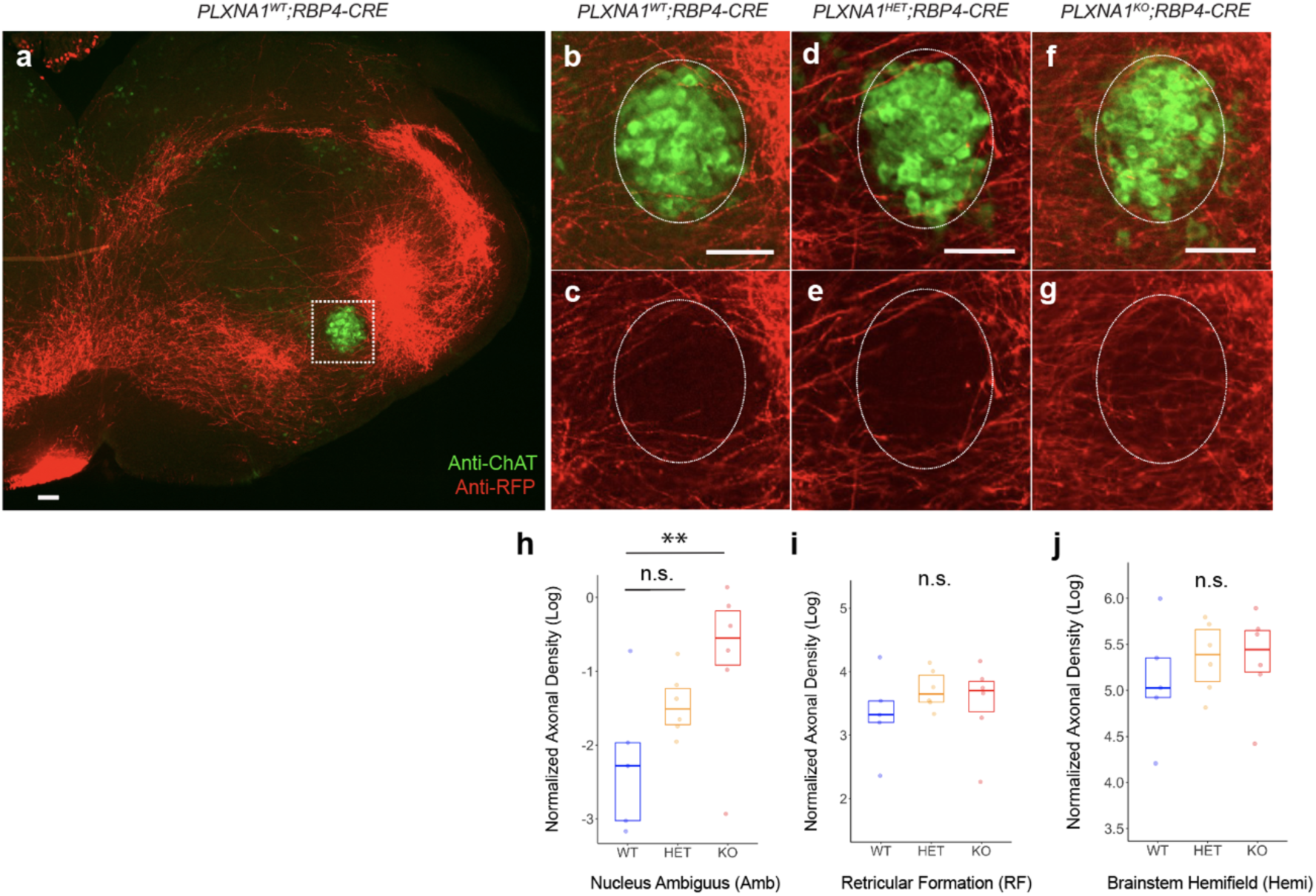
Anterograde tracing of vocal cortico-motoneuronal projections in conditional knockouts of *PLXNA1* pups. **a**-**c**, AAV1.syn.tdTomato injected into primary motor cortex (M1) of P5 *PLXNA1* conditional knockout pups (N_WT_=5, N_HET_=6, N_KO_=5, mice injected) followed by analysis of tdTomato+ (red) cortical projections seven days post-inoculation. tdTomato+ cortical axons are found throughout the brainstem with reduced innervation in the vicinity of ChAT+ (green) motor neurons in the nucleus ambiguus, Amb (dotted box). **d**-**g**, Increased axonal fibers were are found in region of interest (dotted circle) in *PLXNA1^HET^;RBP4-CRE* and *PLXNA1^KO^;RBP4-CRE* mice. **h**-**j**, Axonal density in Amb, reticular formation (RF), and brainstem hemifield (hemi), contralateral to the side of injection, normalized to cortical injection area. **h,** tdTomato+ axonal projections in Amb were significantly higher in *PLXNA1^KO^;RBP4-CRE* mutant pups compared to for *PLXNA1^WT^;RBP4-CRE* controls (p=0.0078, t=-3.42, df=18.8). No significant differences were detected between *PLXNA1^KO^;RBP4-CRE* and *PLXNA1^HET^;RBP4-CRE* (p=0.282, t=-1.57, df=18.8), or *PLXNA1^HET^;RBP4-CRE* and *PLXNA1^WT^;RBP4-CRE* (p=0.160, t=-1.92, df=18.8). **i**,**j**, No significant pairwise differences were observed between *Plxna1* mutant and wildtype pups in the adjacent RF or overall hemifield of the contralateral brainstem relative to the side of AAV1 injection. Overall axonal density varied significantly across brain areas (p<0.001, df=2, F=2110). Linear mixed models were used to predict axonal density using genotype and age as main effect variables and individual mouse pups as random effect variable with repeated measures. Post hoc comparisons used Tukey adjustment for multiple tests. Boxplots visualize the 75^th^ upper and lower 25^th^ quartile of data with horizontal lines indicating median values. Points represent averages of axonal density from 3-5 non-consecutive sections per animal. Scale bars, 100µm. ‘***’ p<0.001, ‘**’ p<0.01, ‘*’ p<0.05, ‘_#_’ p<0.1, ‘n.s.’ non-significant.

**Supplementary Fig. 3.**
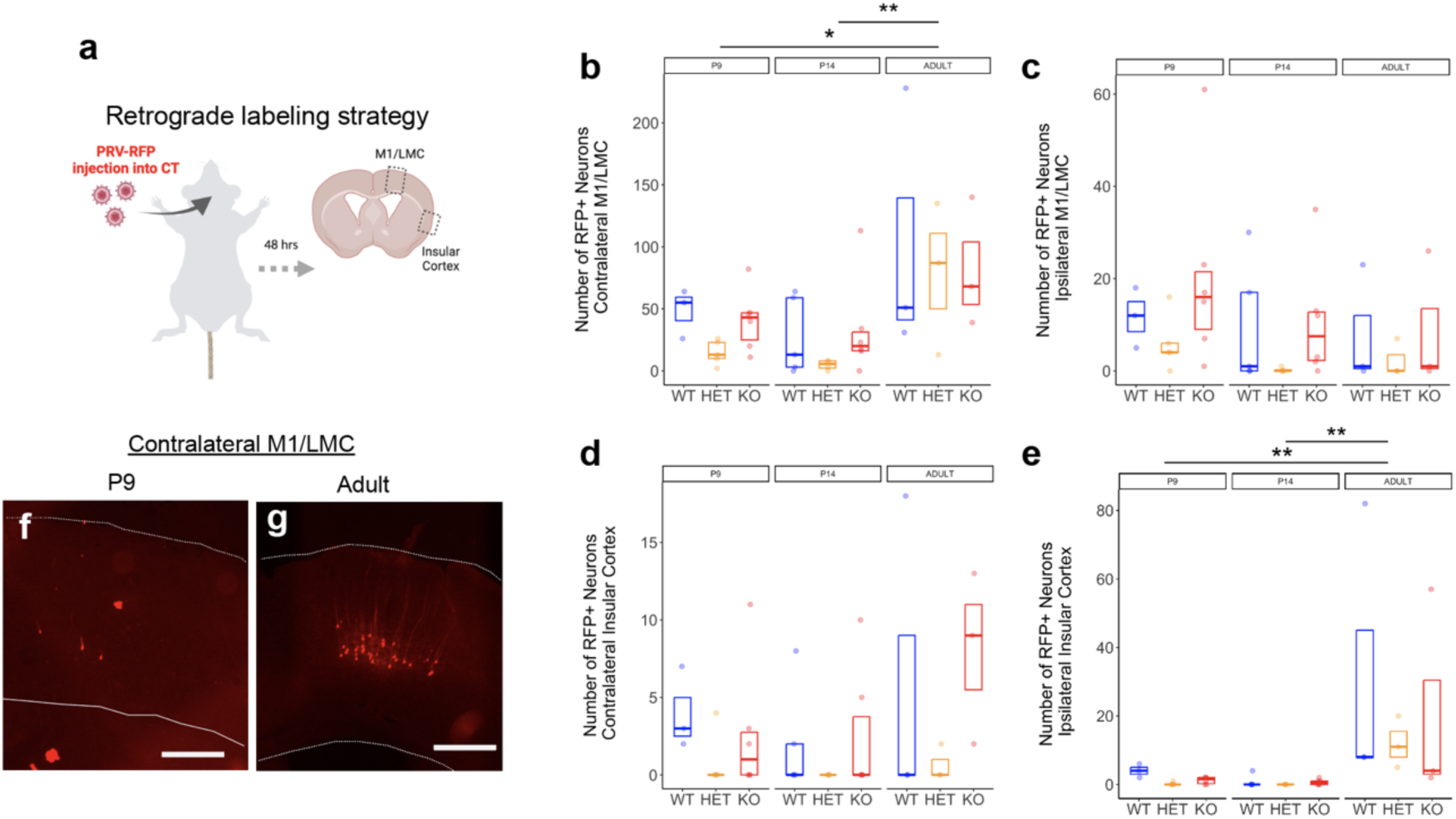
Retrograde labeling of cortical neurons with connectivity to laryngeal muscles. **a**, Pseudo-rabies virus expressing red fluorescent protein (PRV-RFP) was injected into the cricothyroid (CT) muscle of *PLNXA1* conditional knockout mice (P9: N_WT_=3, N_HET_=5, N_KO_=6; P14: N_WT_=5, N_HET_=4, N_KO_=6; Adults: N_WT_=3, N_HET_=3, N_KO_=3). The number of retrogradely labeled RFP+ neurons quantified 48-72 hrs post-inoculation in the deep layers of the primary motor cortex/laryngeal motor cortex (M1/LMC) and insular cortex. Few, if any, RFP+ neurons were observed in more superficial layers of the cortex. **b**, Quantity of RFP+ neurons in the contralateral M1/LMC were not significantly impacted by genotype (p=0.270, df=2, F=1.36) but significantly increased with age (p=0.00168, df=2, F=7.80) with adults having significantly more compared to P9 (p=0.0108) and P14 (p=0.00154) pups. **c**,**d**, Retrogradely RFP+ neurons in the ipsilateral M1/LMC and contralateral insular cortex were not significantly influenced by genotype or age. **e**, Age also had a notable impact on the number of RFP+ neurons in the ipsilateral insular cortex (p=0.00149, df=2, F=7.98) with adults having significantly more labeled neurons than P19 (p=0.005) and P14 (p=0.00199) pups. Boxplots visualize the 75^th^ upper and lower 25^th^ quartile of data with horizontal lines indicating median values. Points represent total number of neuronal counts summed over 5 non-consecutive 50µm sections per animal. **f**,**g**, Example of retrogradely labeled RFP+ neuron in the contralateral M1/LMC in (**f**) P9 pups and (**g**) adult animals. Scale bars, 500µm. Dotted lines indicated basal and apical cortical boundaries. ‘***’ p<0.001, ‘**’ p<0.01, ‘*’ p<0.05, ‘_#_’ p<0.1.

**Supplementary Fig. 4.**
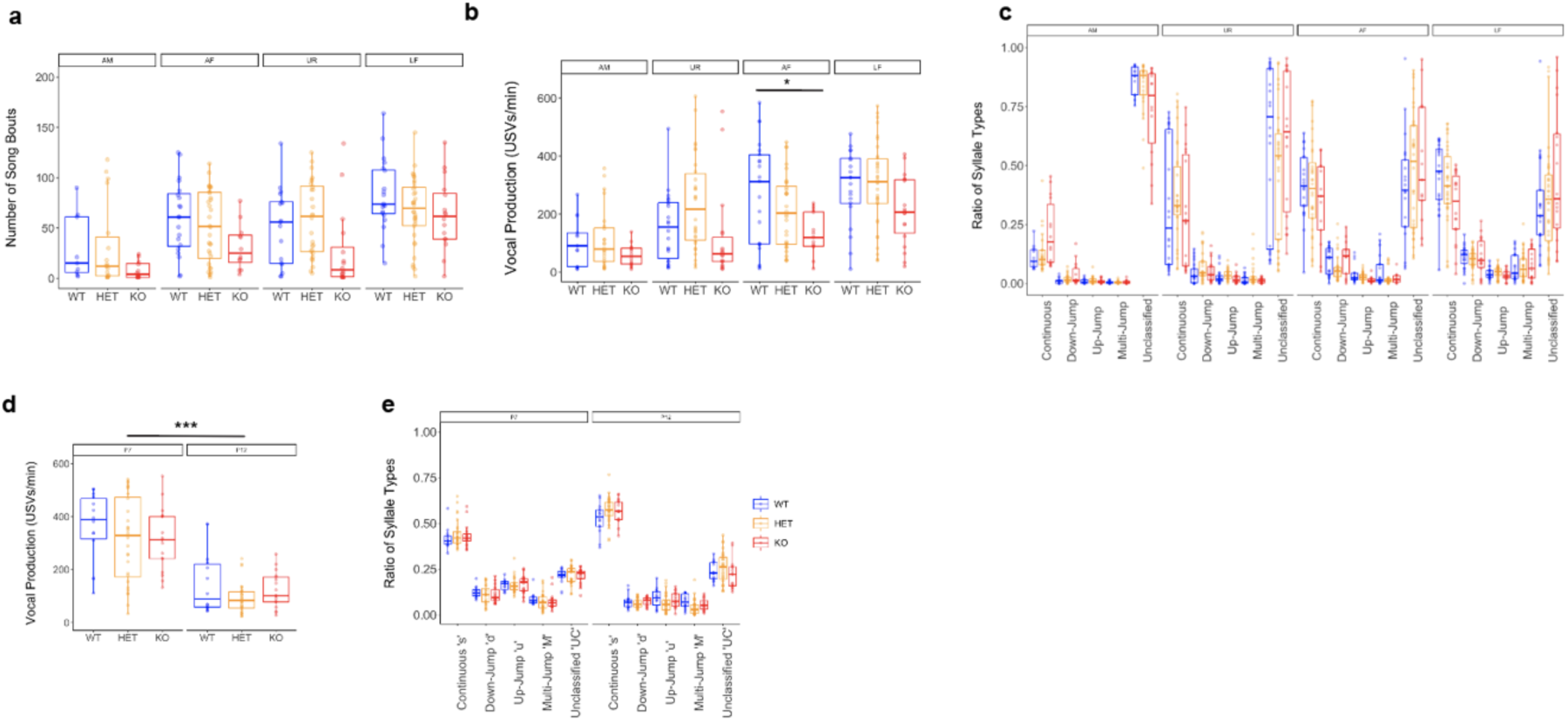
Vocal production in *PLXNA1* conditional knockout pup and adult mice. **a**, Number of song bouts detected from *PLXNA1* conditional knockout adults across anesthetized male (AM), anesthetized female (AF), female urine (UR), and live female (LF) social contexts. **b**, Number of individual USVs produced during each five-minute recording session per animal varied significantly by social context (p=3.27×10-15, df=3, F=27.5) but not genotype (p=0.144, df=2, F=2.11). **c**, Proportion of syllable types, characterized as simple ‘s’, or down-pitch jumps ‘d’, up-jumps ‘u’, multiple pitch jumps ‘M’, or uncategorized ‘UC’, produced during each recording session across social contexts. The ratio of syllable types emitted from adult mice were not impacted by social context (p=1, df=3, F=0) or genotype (p=1, df=2, F=0), but varied across syllable types (p<0001, df=4, F=656), as previously reported^32^. **d**, Production of pup USVs, recorded during five minutes of maternal isolation, decreased from P7 to P12 (p=1.3×10-15, df=1, F=118) with no significant impact of genotype (p=0.205, df=2, F=1.62). **e**, Ratio of syllable types were not impact by age (p=1, df=1, F=0) or genotype (p=1, df=2, F=0), but varied significantly by syllable type (p<0001, df=4, F=1148). Linear mixed models were used to estimate main effects of genotype, age or social context, and syllable type with repeated measures from individual mice modeled as random effects. Post hoc comparisons used Tukey adjustment for multiple test correction. Boxplots visualize the 75^th^ upper and lower 25^th^ quartile of data with horizontal lines indicating median values and whiskers covering the full range of data excluding statistical outliers. ‘***’ p<0.001, ‘**’ p<0.01, ‘*’ p<0.05, ‘_#_’ p<0.1.

**Supplementary Fig. 5.**
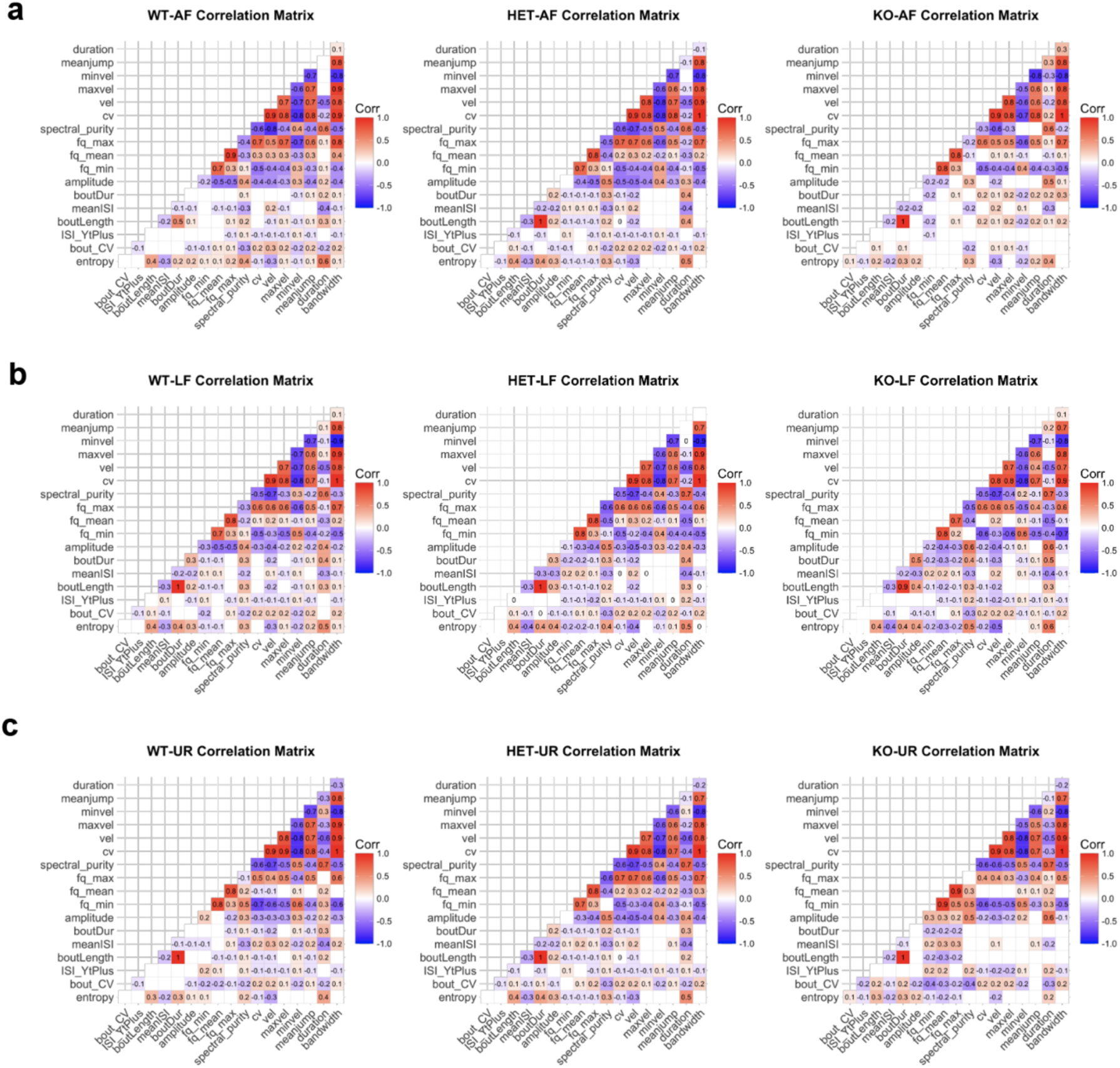
Correlational analysis of song bout features across three social contexts. **a**-**c**, Correlational matrices were computed on song bout features from anesthetized female (AF), female urine (UR), and live female (LF) social contexts using Pearson’s t-distribution product correlation coefficient test. Significant positive (red) and negative (blue) correlations are visualized along with the level of correlation. Similar correlational results and trends across genotypes were obtained using alternative rank-based Spearman correlational tests.

## Supplementary Tables

**Supplementary Table. 1 | Statistical analysis of USVs recorded from adult mice across four social contexts.** ANOVAs were performed on data using the average values of each acoustic feature measured during a recording session of mice in four social contexts: anesthetized male (AM), anesthetized female (AF), female urine (UR), and live female (LF) social contexts. Linear mixed models were generated with genotype, social context, and syllable type as main effect predictors, and individual mice as random variables with repeated measures. The degrees of freedom (df), F values (F), and p values (p) are provided for each main effect variable and respective interactions. ‘***’ p<0.001, ‘**’ p<0.01, ‘*’ p<0.05, ‘_#_’ p<0.1.

**Supplementary Table. 2 | Post hoc analysis of USVs recorded from adult mice across four social contexts.** Post hoc analysis with Tukey adjustment for multiple test correction per acoustic feature was performed using linear mixed models previously generated for ANOVAs investigating vocal behavior across four social contexts: anesthetized male (AM), anesthetized female (AF), female urine (UR), and live female (LF) social contexts. Syllable types include simple ‘s’, down-pitch jumps ‘d’, up-jumps ‘u’, multiple pitch jumps ‘M’, or uncategorized ‘UC’, The estimate, standard error (SE), degrees of freedom (df), t-value (t), and p-value (p) are provided for each comparison. ‘***’ p<0.001, ‘**’ p<0.01, ‘*’ p<0.05.

**Supplementary Table. 3 | Statistical analysis of USVs recorded from mouse pups at two stages of development.** ANOVAs were performed on data using the average values of each acoustic feature measured during a recording session in two developmental stages: post-natal day 7 (P7) and P12. Linear mixed models were generated with genotype, age, and syllable type as main effect predictors, and individual mice as random variables with repeated measures. The degrees of freedom (df), F values (F), and p values (p) are provided for each main effect variable and respective interactions. ‘***’ p<0.001, ‘**’ p<0.01, ‘*’ p<0.05, ‘_#_’ p<0.1.

**Supplementary Table. 4 | Post hoc analysis of USVs recorded from mouse pups at two stages of development.** Post hoc analysis with Tukey adjustment for multiple test correction per acoustic feature was performed using linear mixed models previously generated for ANOVAs investigating vocal behavior at two developmental stages: post-natal day 7 (P7) and P12. Syllable types included: simple ‘s’, or having down-pitch jumps ‘d’, up-jumps ‘u’, multiple pitch jumps ‘M’, or uncategorized ‘UC’, The estimate, standard error (SE), degrees of freedom (df), t-value (t), and p-value (p) are provided for each comparison. ‘***’ p<0.001, ‘**’ p<0.01, ‘*’ p<0.05.

## Notes

### Competing Interest Statement

The authors have declared no competing interest.

### Summary of Updates

Figure legends updated. Author ORCID added. Minor text revisions.

